# Acute kidney injury triggers hypoxemia by inducing intravascular neutrophil retention that reduces lung capillary blood flow

**DOI:** 10.1101/2024.02.27.582396

**Authors:** Yohei Komaru, Liang Ning, Carine Lama, Anusha Suresh, Eirini Kefaloyianni, Mark J. Miller, Shinichi Kawana, Hailey M. Shepherd, Wenjun Li, Daniel Kreisel, Andreas Herrlich

## Abstract

Sterile acute kidney injury (AKI) is common in the clinic and frequently associated with hypoxemia that does not improve with dialysis and remains incompletely understood. AKI induces remote lung inflammation with neutrophil recruitment in mice and humans, but which cellular cues establish neutrophilic inflammation and how it contributes to hypoxemia is not known. Here we report that AKI induces rapid intravascular neutrophil retention in lung alveolar capillaries without any significant extravasation into tissue or alveoli, causing hypoxemia by reducing lung capillary blood flow in the absence of any significant lung interstitial or alveolar edema. In contrast to direct ischemic lung injury, lung neutrophil recruitment during remote lung inflammation did not require cues from intravascular non-classical monocytes or tissue-resident alveolar macrophages. Instead, lung neutrophil retention depended on neutrophil chemoattractant CXCL2 released by activated intravascular classical monocytes. Comparative single-cell RNA-sequencing analysis of direct and remote lung inflammation revealed that alveolar macrophages are highly activated and produce the neutrophil chemoattractant CXCL2 only in direct lung inflammation. Establishing a CXCL2 gradient into the alveolus by intratracheal administration of CXCL2 during AKI-induced remote lung inflammation enabled neutrophils to extravasate. We thus discovered important differences in lung neutrophil recruitment in direct versus remote lung inflammation and identified lung capillary neutrophil retention that negatively affects oxygenation by causing a ventilation-perfusion mismatch as a novel driver of AKI-induced hypoxemia.

## Introduction

Sterile AKI is a common clinical condition that carries high mortality, particularly when it occurs with respiratory failure (1–3). AKI-associated respiratory failure is incompletely understood and thought to result from lung interstitial and alveolar edema as a result of endothelial leakage in response to systemic inflammation induced by AKI (4, 5). In mice (4, 6–10) and humans (11, 12), AKI-induced systemic inflammatory mediators cause remote lung inflammation with neutrophil accumulation, but little to no alveolar inflammatory or serous exudates (reviewed in (4, 7, 8)). We showed for the first time that mice are hypoxemic 24 hours after AKI despite the lack of significant alveolar edema or exudates and that prevention of lung neutrophil accumulation prevented hypoxemia (6). Which cellular cues attract neutrophils into the lung during remote inflammation and how neutrophils contribute to hypoxemia after AKI, however, remains largely unresolved.

Here we show that during remote lung inflammation after AKI the vast majority of neutrophils do not extravasate. Intravascular retention of neutrophils in pulmonary capillaries obstructs lung capillary blood flow and significantly reduces oxygenation by 2–6 hours after AKI in the absence of any significant interstitial or alveolar edema or inflammatory exudates. We show that in remote lung inflammation, unlike in direct lung injury, non-classical monocytes and alveolar macrophages are not required to attract neutrophils to the lung and, instead, implicate the neutrophil chemoattractant CXCL2 released by intravascular classical monocytes in capillary neutrophil retention. Comparative single-cell RNA-seq analysis of direct and remote lung inflammation revealed that alveolar macrophages produce CXCL2 only during direct lung inflammation. Establishing a CXCL2 gradient into the alveolus by intratracheal administration of CXCL2 during AKI-induced remote lung inflammation enabled neutrophils to extravasate. We thus describe important differences in lung neutrophil recruitment in direct versus remote lung inflammation. We have uncovered a novel mechanism that underlies AKI-induced hypoxemia, namely intravascular neutrophilic inflammation with capillary perfusion deficits that worsen oxygenation despite proper alveolar function and ventilation. We hypothesize that lung intravascular neutrophil retention after AKI may protect the organism from overt systemic action of neutrophils and inflammatory mediators by sequestering them in the lung vasculature, yet at the risk of hypoxemia due to reduced lung alveolar capillary perfusion and oxygen uptake. Our findings impact the understanding of AKI-associated respiratory failure in general and also may help explain the negative impact AKI has on respiratory failure and mortality in patients with acute respiratory distress syndrome (ARDS) (13).

## RESULTS

### Neutrophils recruited to the lung after AKI do not extravasate from lung vessels

During direct tissue injury neutrophils typically establish inflammation by extravasating from the circulation into the injured tissue (14, 15). During remote lung inflammation after AKI we and others have previously found that few if any neutrophils are detectable in bronchoalveolar lavage fluid (reviewed in (4, 5, 16), indicating that they do not transmigrate through the pulmonary epithelial barrier. However, whether neutrophils exit the vascular space during remote lung inflammation is not well understood. We induced bilateral renal ischemia-reperfusion injury (IRI) in mice as a model of severe AKI. In wildtype (*wt*) mice, blood urea nitrogen (BUN) levels typically reach 100–150 mg/dL 24 hours after IRI (20-minute ischemia time) (Figure 1A). Flow cytometry analysis confirmed a significant increase in the total number of lung neutrophils after AKI, irrespective of whether the lungs were perfused with phosphate-buffered saline (PBS) at the time of sample preparation (Figure 1B). This suggested that neutrophils either extravasated into the lung interstitium or that they could not be flushed out due to strong vascular adhesion or vascular obstruction. To determine the localization of neutrophils, we injected an APC-labeled anti-Ly6G antibody intravenously 10 minutes prior to sacrifice and a BV421-labeled anti-Ly6G antibody was added to lung single cell suspensions generated from the same animal after sacrifice (17). Only intravascular neutrophils are double positive (APC^+^/BV421^+^), while extravasated neutrophils are single positive (BV421^+^) (Figure 1C). We used direct lung injury by intratracheal instillation of lipopolysaccharide (LPS) as a positive control for neutrophil extravasation. 24 hours after bilateral kidney IRI to induce AKI (ischemic AKI), the total number of neutrophils in the lung significantly increased over sham levels and was comparable to the increase observed after direct lung injury with LPS (Figure 1D). However, after AKI >99.5% of neutrophils remained within the intravascular compartment in contrast to direct lung injury where >80% of neutrophils had extravasated (Figure 1E–F). Sterile kidney tissue injury-recruited lung neutrophils may therefore impact oxygenation by impeding blood flow in lung vessels. This is consistent with our published findings that AKI-induced remote lung inflammation with hypoxemia in our model lacks significant alveolar edema or inflammatory exudates (6).

**Figure 1.**
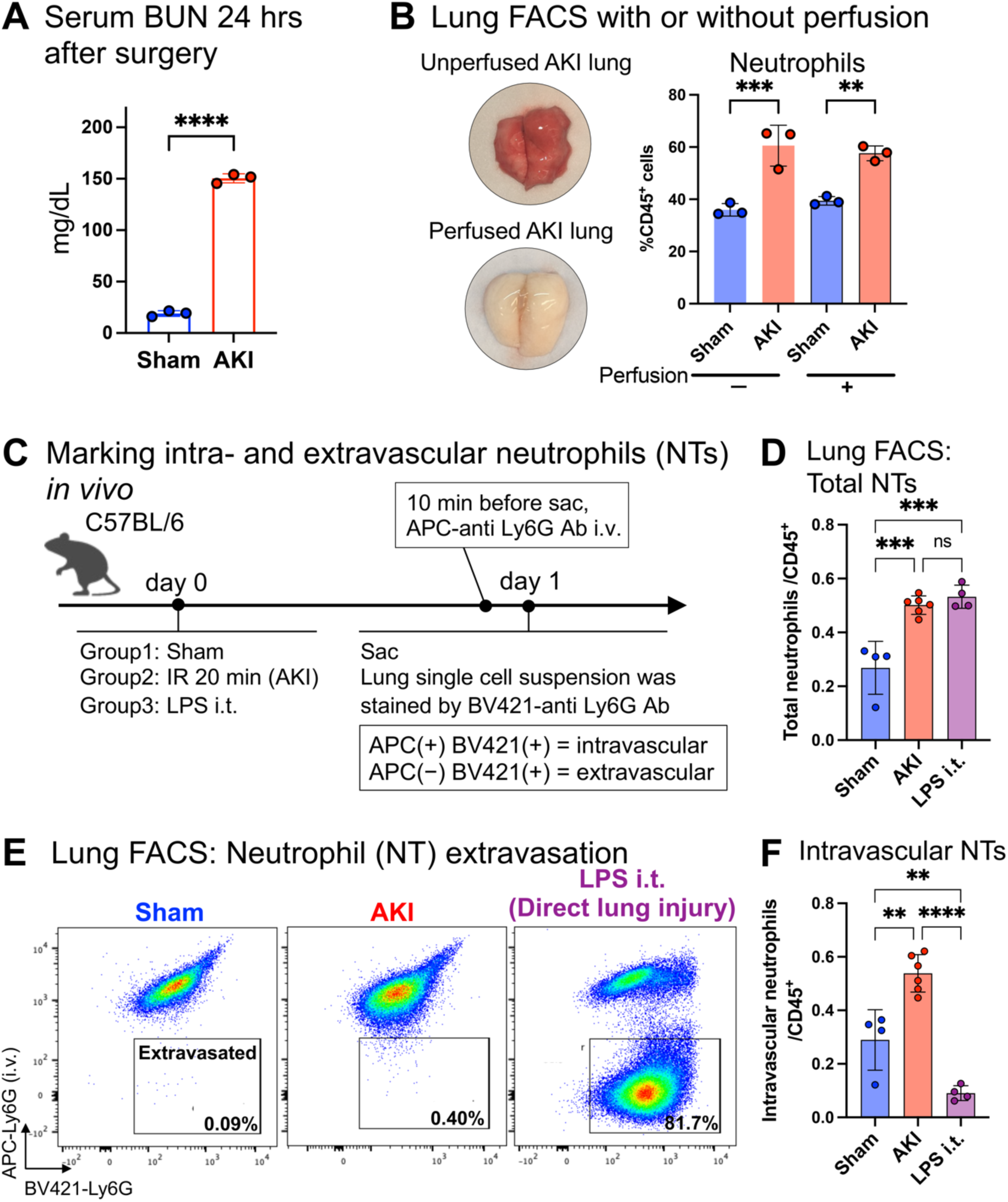
AKI induces intravascular neutrophil accumulation in the lung. (**A**) Serum blood urea nitrogen (BUN) levels 24 hours after sham or AKI in wildtype mice. (**B**) Representative image of mouse lung before and after perfusion with phosphate-buffered saline (PBS). Total neutrophil counts in sham vs. AKI lungs were quantified by flow cytometry (FACS) with or without prior lung perfusion. Neutrophils/CD45^+^ cells: unperfused, 36.0% vs. 60.5%; perfused, 39.4% vs. 57.6%; in both sham vs. AKI, p < 0.01. (**C**) Schematic of the in vivo neutrophil extravasation assay using two different anti-Ly6G antibodies coupled to different fluorescent labels (APC or BV421). (**D**) FACS quantification of lung total neutrophils after sham, AKI (remote lung inflammation), or intratracheal (i.t.) lipopolysaccharide (LPS) (direct lung injury). (**E**) Identification of extravasated neutrophils by FACS. Ly6G fluorescence (APC over BV421) is analyzed in CD45^+^, CD11b^+^, Gr1^high^ cells in sham animals (left panel), AKI animals (middle), and animals subjected to direct lung injury by LPS i.t. (left); single positive cells (BV421) are extravascular, double positive cells (APC/BV421) are intravascular. (**F**) FACS quantification of intravascular lung neutrophils (after PBS perfusion) for sham, AKI, or LPS i.t.. Data in (B, D, F) represent mean ± standard deviation (SD) and n = 3 to 6 mice per group. ns: not significant. **P < 0.01, ***P < 0.001, ****P < 0.0001

### Intravascular lung capillary neutrophil retention causes hypoxemia by reducing lung capillary perfusion

To determine early events that establish remote lung inflammation in real time, we investigated monocyte and neutrophil behavior in lungs using intravital two-photon microscopy at two hours after AKI as compared to uninjured sham controls. We used *Ccr2^gfp/+^* mice that allow in vivo tracking of classical monocytes via GFP fluorescence (green) (18). Neutrophils, blood flow and vessels were visualized in vivo by i.v. injection of a fluorescently labeled anti-Ly6G antibody (red), 1-micron fluorescent microbeads (white), and Qtracker655 (violet), respectively. Two hours after sham operation few neutrophils, very few static microbeads, and few classical monocytes were detected in the lung. Classical CCR2^+^ monocytes were largely localized at a significant distance from neutrophils (Movie S1). By contrast, two hours after AKI, we observed a large number of neutrophils lining up in lung capillaries that did not extravasate and formed neutrophil “trains”, which were generally tightly associated with at least one classical monocyte “locomotive” (see inset on right) (Figure 2A and Movie S2). Intravascular neutrophil speed was decreased to levels indicative of crawling/arrest (Figure 2B) and the presence of a large number of immobile intravascular 1-micron beads after AKI suggested severely reduced blood flow in the lung capillary microcirculation (Figure 2C). As additional supportive evidence for reduced capillary blood flow, we detected evidence of lung micro-thrombosis by anti-platelet (CD41a) and anti-fibrinogen immunofluorescence co-staining (about 6% of the lung area), in AKI but not in sham lungs (Supplemental Figure 1). The average distance between classical monocytes and neutrophils was significantly reduced after AKI, raising the possibility of chemoattraction (Figure 2D). To detect neutrophil train formation and monocyte/neutrophil behavior at even earlier time points we obtained 3D timelapse images of mouse lungs immediately after AKI (5–10min after reperfusion) (Figure 2E, and Movie S3). Already at this very early time point we observed an increased number of neutrophils and CCR2^+^ classical monocytes in the lung. Overall monocyte speed was significantly reduced, with many monocytes not moving at all. At the same time many neutrophils were already present in the lung moving at much higher speeds than monocytes. In some instances, we could already observe the formation of “neutrophil trains” capped by a monocyte in lung capillaries. In such examples, neutrophils were moving toward an immobile monocyte, suggesting that the monocyte may send a neutrophil chemoattractant signal (Figure 2E). Of note, this intravascular neutrophil retention phenotype after AKI is very different from neutrophil behavior in direct lung injury due for example to warm lung ischemia-reperfusion injury in a murine syngeneic lung transplant model (19). In direct lung injury neutrophils do not remain in the intravascular space to form trains, but instead extravasate and form large alveolar swarms as early as 2 hours after reperfusion (Video S4, Q-dot vascular dye: *red*; neutrophils: *green*). Taken together, our findings indicate that after AKI neutrophils accumulate very rapidly in lung capillaries where they assemble into blood flow impeding “trains”, possibly involving chemoattraction between neutrophils and monocytes.

**Figure 2.**
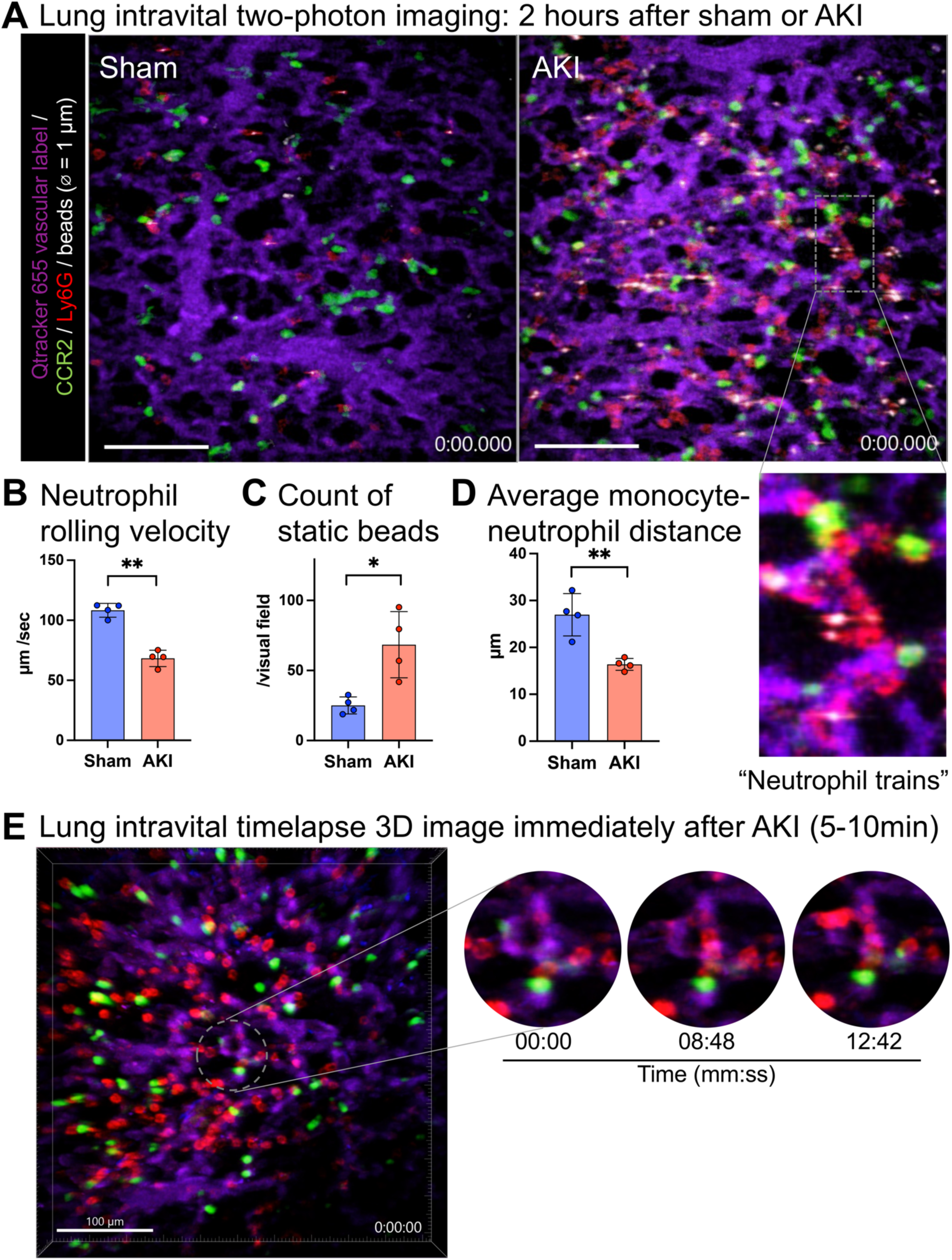
Intravascular lung capillary neutrophil train formation after AKI. (**A**) Intravital two-photon imaging of sham or AKI lungs 2 hours after AKI using *Ccr2^gfp/+^* mice and in vivo staining of neutrophils; CCR2^+^ monocytes (green), Ly6G^+^ neutrophils (red). Blood flow is assessed with 1-micron beads (white) and lung capillary circulation is labeled with intravenous injection of quantum dots (purple). Magnified inset image shows a “neutrophil train” in a lung capillary, illustrating vessel capillary circulation is labeled with intravenous injection of quantum dots (purple). Magnified inset image shows a “neutrophil train” in a lung capillary, illustrating vessel occlusive accumulation of neutrophil “trains” attached to CCR2^+^ monocyte “locomotives”. Full videos are available in supplementary materials. Scale bar = 100 µm. (**B**–**D**) Quantification of intravital imaging videos: speed of neutrophil rolling (B), the number of static beads (C), and the averaged distance between CCR2^+^ monocytes and neutrophils (D). (**E**) Lung intravital timelapse 3D image immediately after AKI (5–10 min). Magnified images on the right illustrate the process of neutrophil (red) train formation in the presence of a CCR2^+^ monocyte (green). Full timelapse video is available as Movie S3. Scale bar = 100 µm. Data in (B-E) represent mean ± SD and n = 4 to 6 mice per group. *P < 0.05, **P < 0.01, ***P < 0.001 compared to sham.

We next asked whether the observed significant reduction in lung capillary blood flow could reduce blood oxygenation in the absence of significant interstitial/alveolar edema or alveolar exudates (6). Compared to control conditions, arterial blood gas measurements revealed that, while no significant oxygenation defect was present at 2 hours, hypoxemia developed by 6 hours after AKI and correlated with the degree of kidney injury (Supplemental Figure 2). Oxygenation was reduced by ∼16% at 6 hours (Figure 3A), similar to our previous observations at 24 hours after AKI in this model (6). No significant alveolar edema or exudates and only small alveolar wall thickness increases of 10–15% were detectable at 2 and 6 hours after AKI (Figure 3B). Importantly, hypoxemia occurred between 2–6 hours after AKI without any substantial changes in alveolar wall thickness during the same time frame, implicating impaired capillary perfusion resulting in ventilation-perfusion deficits rather than interstitial edema as a cause of hypoxemia. Our findings are consistent with our previous report of only small alveolar wall thickness increases and absence of alveolar edema or exudates up to 24 hours after AKI, and with our previous finding that prevention of neutrophil accumulation during this time frame prevented hypoxemia (6). To further confirm that lung neutrophil accumulation is indispensable for impaired lung capillary perfusion and hypoxemia, we analyzed AKI animals following neutrophil depletion. Pretreatment with anti-Ly6G antibody effectively reduced the number of circulating neutrophils after AKI (Figure 3C), and neutrophil-depleted mice were protected from hypoxemia 6 hours after AKI (Figure 3D). The number of lung fluorescent microbeads intravenously injected 10 minutes before sacrifice was significantly higher in mice without neutrophil depletion (control IgG) than with neutrophil depletion (anti-Ly6G), suggesting that neutrophil retention in the lung is indeed critical for the reduction in capillary flow and hypoxemia (Figure 3E).

**Figure 3.**
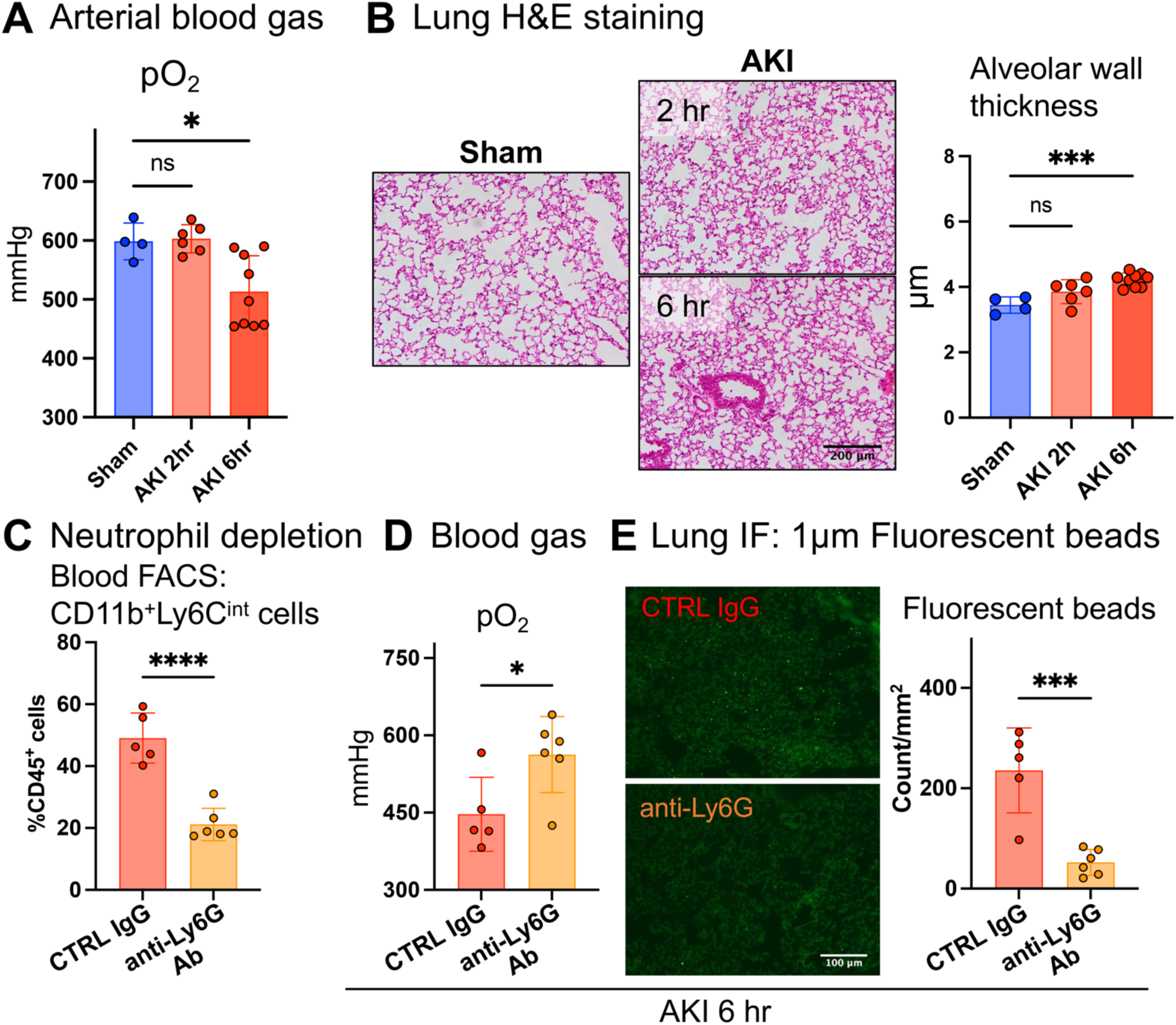
Hypoxemia is observed early after AKI in the absence of impaired ventilation or overt lung edema. (**A**) Arterial blood gas analysis after sham or 2–6 hours after AKI. Arterial blood samples were collected directly from ascending aorta under mechanical ventilation using 100% O_2_. (**B**) Lung hematoxylin and eosin (H&E) staining with alveolar wall thickness measurements. Scale bar = 200 µm. (**C**–**E**) Neutrophil depletion experiment. (C) Quantification of circulating neutrophils (CD11b^+^Ly6C^int^ cells) following two injections of anti-Ly6G antibody. Ly6G was not used as a neutrophil marker in FACS analysis due to epitope protection by the Ly6G antibody injected for neutrophil depletion. (D) Arterial blood gas analysis 6 hours after AKI. (E) Quantification of lung fluorescent beads injected 10 minutes prior to sacrifice as a surrogate marker for stagnation of lung microcirculation. Data represent mean ± SD. *P < 0.05, ***P < 0.001, ****P < 0.0001, compared to sham or control.

### Rapid lung capillary neutrophil capture is enhanced by decreased neutrophil deformability but not classical neutrophil-endothelial cell interactions

Circulating neutrophils are attracted to sites of inflammation based on identifiable sequential interactions of different neutrophil cell surface ligands with endothelial cell surface receptors, encompassing canonical steps of neutrophil capture, rolling, crawling, arrest, and migration, prior to potential extravasation (20). Whether neutrophil-endothelial interactions are important in lung neutrophil recruitment after AKI is unknown. Based on our observations of neutrophil crawling/arrest during intravital imaging already 2 hours after AKI, we tested blockade of key neutrophil-endothelial cell interactions that mediate neutrophil crawling/arrest and migration. The integrins LFA-1 (CD11a/CD18) and Mac-1 (CD11b/CD18) play overlapping roles in neutrophil arrest and migration (20, 21). Supporting the potential involvement of these integrins, in an analysis of single-cell RNA sequencing (scRNA-seq) datasets in sham and AKI mice ICAM-1, a ligand for LFA-1 or Mac-1, was found upregulated in lung endothelium on day 1 after AKI (Figure 4A). Supplemental Figure 3 contains a detailed description of our analysis using our previously published dataset (6) together with additional newly generated lung scRNA-seq data. Despite endothelial upregulation of ICAM-1, blockade of CD18 (Figure 4B) or LFA-1 (Supplemental Figure 4) did not reduce early neutrophil recruitment into the lung at two hours after AKI, suggesting that neutrophil accumulation during remote lung inflammation does not follow the classical neutrophil recruitment paradigm delineated largely in models of direct tissue injury (22).

**Figure 4.**
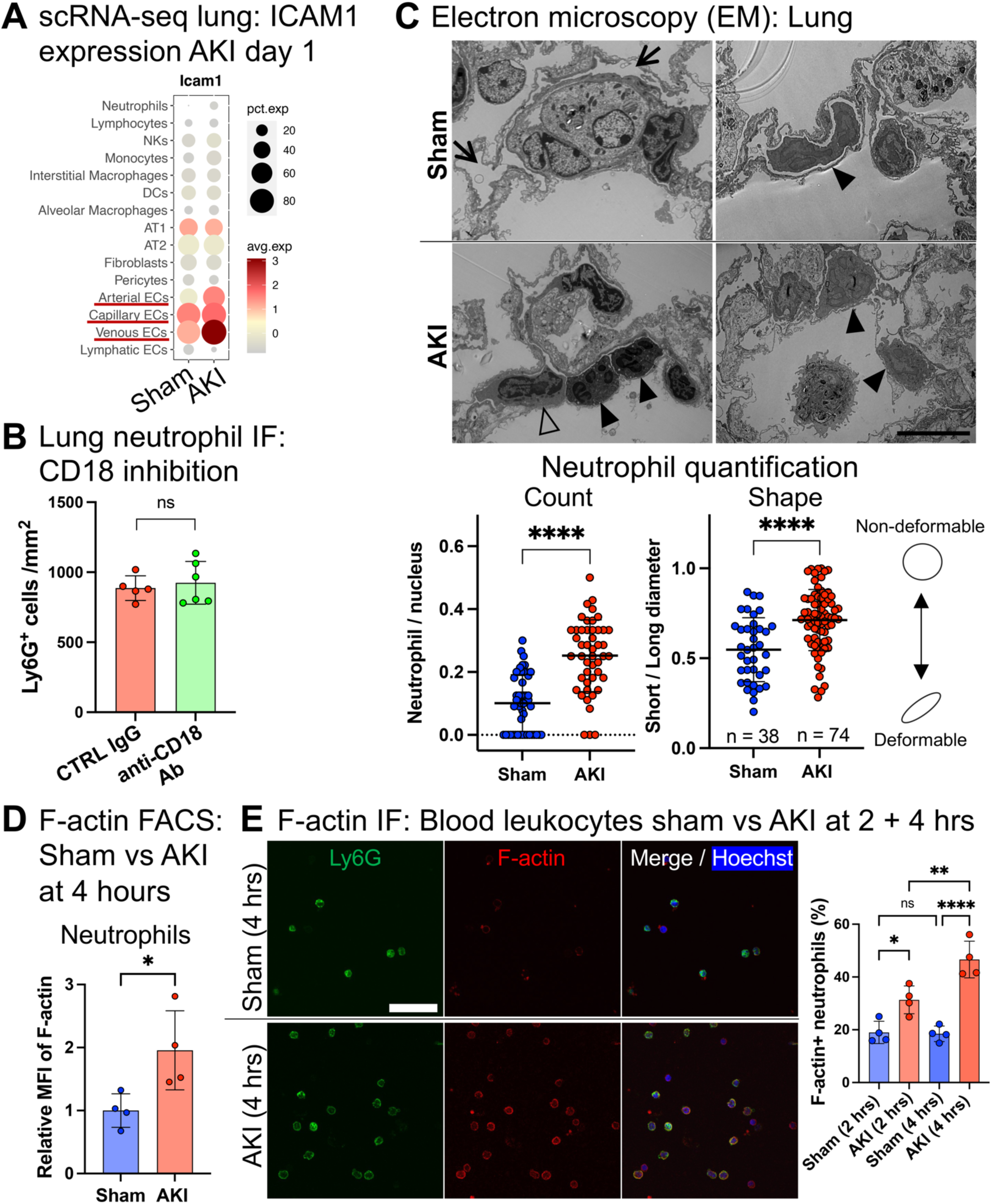
Rapid lung capillary neutrophil capture is enhanced by decreased neutrophil deformability but not classical neutrophil-endothelial cell interactions. (**A**) Single-cell RNA seq analysis (see also Supplemental Figure 2): Expression of adhesion molecule, ICAM1, in lung cell types after sham or AKI. The dot size denotes percentage of cells expressing each gene, and the color scale represents average gene expression levels. (**B**) Anti-CD18 antibody blockade vs IgG control in C57BL/6 mice: Quantification of lung neutrophils by Ly6G immunofluorescence staining after AKI. (**C**) Representative images of sham vs. AKI lung obtained by electron microscopy (EM) with quantification (graphs). Arrows indicate empty vessel lumina, while full arrowheads point out deformable neutrophils in sham or non-deformable neutrophils in AKI lung; in the AKI lung an open arrowhead is pointing out a monocyte “locomotive” leading a non-deformable neutrophil train. Right lower image shows crawling or arrested neutrophils (closed arrowheads). EM images were obtained from 3 mice/group for sham and AKI. Scale bar = 10 µm. Left graph: neutrophil count is presented as number of neutrophils per total nuclei in each image. Right graph: the ratio of short over long diameter in each neutrophil is shown as an indicator of its morphology. Short/long ratio closer to 1.0 is indicative of a more rounded non-deformable shape. (**D**) Quantification of F-actin in circulating neutrophils by flow cytometry. Using fluorescently-labeled phalloidin, geometric mean fluorescence intensity (MFI) is analyzed and compared between sham and AKI. (**E**) F-actin staining of leukocytes isolated from circulating blood 2 and 4 hours after sham or AKI. Scale bar = 40 µm. Percentage of F-actin polymerized Ly6G^+^ cells (neutrophils) are quantified in graph. n = 4 to 6 mice/group. Data represent mean ± SD. ns: not significant, *P < 0.05, **P < 0.01, ****P < 0.0001

We next hypothesized that rapid neutrophil capture in lung capillaries after sterile kidney tissue injury (<1 hour as published by us in (6) and beginning already at 5–10min after AKI Figure 2 E) may not depend on a specific capture signal that emanates from the lung. Instead, it could depend on reported changes in cellular deformability of primed neutrophils (23) stimulated by circulating inflammatory mediators, as has been described in humans with systemic inflammatory response due to sepsis, trauma, or direct lung injury (24, 25). This is relevant because neutrophils have to be deformable to pass through the pulmonary microcirculation as the average capillary diameter is approximately 7.5 microns (21, 26, 27), significantly smaller than neutrophils and monocytes, which have an average diameter of 9–15 microns (21). Non-deformability of primed neutrophils is conferred by rapid changes in their cellular shape induced by cytoskeletal rearrangement with the hallmark formation of polymerized F-actin bands in sub-cell membrane regions (28). We therefore examined cell shape and presence of F-actin bands in neutrophils after AKI as compared to sham conditions with electron microscopy (EM). This analysis confirmed the intravascular neutrophil accumulation seen with intravital imaging. Neutrophilic cell shape was determined by measuring and comparing the ratio of the longest short to the longest long axis. At 24 hours after AKI, we detected mostly empty vessel lumina (arrows) and few neutrophils (full arrowhead) in sham lungs (Figure 4C, upper images and graph). Most neutrophils had a deformable elongated cellular shape with a smooth cellular surface and no obvious interactions with the endothelium. Mice with AKI showed an increased number of neutrophils within the vasculature (full arrowheads) that exhibited a less or non-deformable rounded shape (Figure 4C, lower images and graph). As seen during intravital imaging, EM revealed monocytes (open arrowhead) in close association with “neutrophil trains” (closed arrowheads) (Figure 4C left lower image). Next, we visualized F-actin polymerization and peripheralization using phalloidin staining in circulating neutrophils isolated from sham or AKI mice. Flow cytometry detected significantly increased F-actin fluorescence in neutrophils 4 hours after AKI compared to sham (Figure 4D). By immunofluorescence staining, 20% of circulating neutrophils isolated from sham animals showed sub-cell membrane F-actin bands, whereas in AKI animals sub-cell membrane F-actin bands were detected in 30% of neutrophils at 2 hours, 40–50% at 4 hours (Figure 4E) and 50% on day 1 after AKI (Supplemental Figure 5). Thus, the accumulation of neutrophils in lungs after AKI is likely significantly promoted by their priming/activation through kidney injury-released circulating factors that reduce their deformability, causing their rapid capture in pulmonary capillary vessels.

### Non-classical monocytes or alveolar macrophages do not drive lung capillary neutrophil retention after AKI

We next aimed to elucidate which cellular cues are needed for AKI-induced lung neutrophil accumulation. In direct lung injury, neutrophil recruitment and extravasation is driven by cues derived from non-classical monocytes (29), alveolar macrophages (30), and classical monocytes (31), respectively. *Nr4a1^-/-^*mice (32) lack non-classical monocytes but not classical monocytes, as shown by Ly6C flow cytometry of circulating monocytes. Classical monocytes in *Nr4a1^-/-^*mice are in fact a little elevated because they cannot transition into non-classical monocytes (Figure 5A). *Nr4a1*^-/-^ mice experienced similar kidney injury in response to bilateral IRI as did *wt* controls (Figure 5B) but showed no differences in lung neutrophil accumulation after AKI (Figure 5C). We next eliminated alveolar macrophages with intratracheal diphtheria toxin (DTX) application in *Cd169^DTR/+^* mice (Figure 5D Experimental Scheme and Supplemental Figure 6). *Cd169^DTR/+^* mice treated with DTX showed comparable kidney injury to *wt* controls (Figure 5E), but alveolar macrophage depletion did not affect lung neutrophil accumulation after AKI (Figure 5F). Thus, non-classical monocytes and alveolar macrophages do not play critical roles in mediating neutrophil accumulation in lungs after AKI.

**Figure 5.**
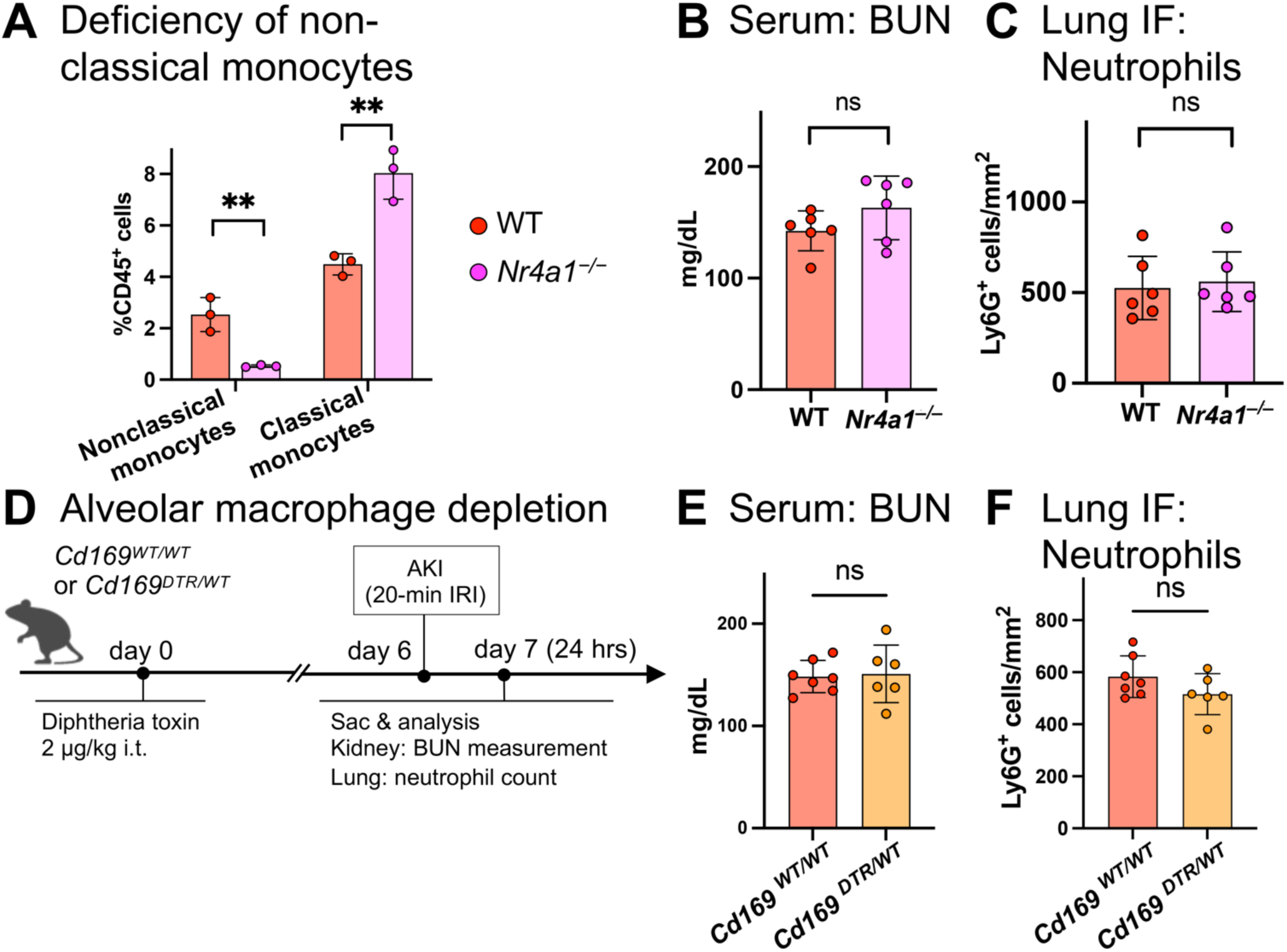
Neither non-classical monocytes nor alveolar macrophages drive capillary neutrophil retention. (**A**) Quantification of blood monocytes in wildtype (*wt*) vs. NR4A1 knockout mice (*Nr4a1^-/-^*). (**B**) Serum BUN levels 24 hours after AKI in wildtype and NR4A1 knockout mice. (**C**) Quantification of lung neutrophils after AKI measured by immunofluorescence staining. (**D**) Schematic of in vivo alveolar macrophage depletion model using diphtheria toxin and CD169-DTR heterozygous mice. (**E**) Serum BUN levels 24 hours after AKI in CD169-DTR heterozygous mice and their littermate controls. (**F**) Quantification of lung neutrophils after AKI measured by immunofluorescence staining in CD169-DTR heterozygous mice and their littermate controls. Data represent mean ± SD. ns: not significant, **P < 0.01.

### CCR2^+^ classical monocytes and CXCL2 drive lung capillary neutrophil retention

Since classical monocytes appeared to interact with neutrophils based on intravital imaging and EM in lung capillaries, we examined their role in establishing lung capillary neutrophil trains. First, we selectively depleted CCR2^+^ classical monocytes by administration of anti-CCR2 antibody (33) (Figure 6A). This antibody does not deplete circulating neutrophils (34). Anti-CCR2 antibody pre-treatment depleted classical (CD11b^+^, Ly6G^−^, CCR2^+^, Ly6C^+^), but not non-classical monocytes (CD11b^+^, Ly6G^−^, Ly6C^−^) (Figure 6B). No differences in kidney injury between anti-CCR2 or control-antibody treated animals were observed (Figure 6C). However, depletion of classical monocytes inhibited lung neutrophil and also interstitial macrophage accumulation after AKI almost completely as compared to control (Figure 6D). These results identify classical CCR2^+^ monocytes as key upstream players in AKI-induced lung neutrophil accumulation.

**Figure 6.**
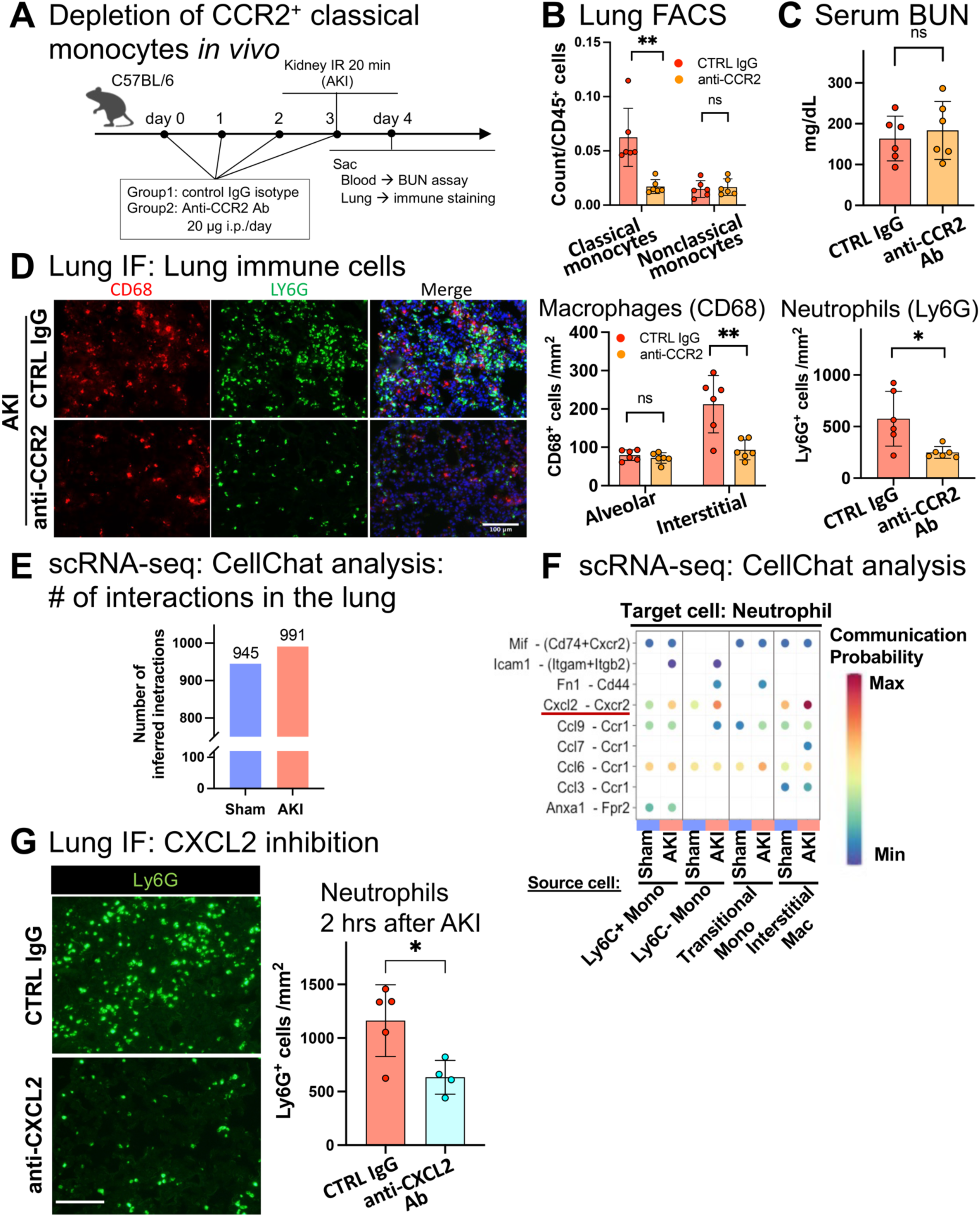
CCR2^+^ classical monocytes and CXCL2-CXCR2 signaling drive lung capillary neutrophil retention after AKI. (**A**) Schematic of classical monocyte depletion using anti-CCR2 antibody in vivo. (**B**) Lung monocytes after pretreatment with anti-CCR2 antibody or control IgG. (**C**) Serum BUN: Kidney injury level on day 1 after AKI. (**D**) Lung immunofluorescence staining after AKI in control vs. anti-CCR2 antibody-treated animals: alveolar and interstitial macrophages (CD68^+^, red) and neutrophils (Ly6G^+^, green). Hoechst 33342 dye (blue) was used to visualize nuclei. Scale bar = 100 µm. (**E**) Total number of cell-cell communications inferred by CellChat based on scRNA-seq analysis in sham vs. AKI (also see Supplemental Figure 2). (**F**) Cell-cell communications from monocytes or macrophages to neutrophils were predicted at single ligand-receptor resolution (unbiased); the neutrophil chemoattractant CXCL2 in monocytes/macrophages and its receptor CXCR2 in neutrophils were predicted as significantly increased in AKI vs. sham and are underlined. (**G**) Lung immunofluorescence staining for Ly6G^+^ neutrophils (green) in wildtype C57BL/6 mice injected with anti-CXCL2 or control antibody and subjected to AKI. Scale bar = 100 µm. Data represent mean ± SD and n = 4 to 6 mice per group. ns: not significant, *P < 0.05, **P < 0.01

To identify possible immune cell signals that recruit neutrophils, in particular emanating from classical CCR2^+^ monocytes, we performed cell-cell communication analysis with CellChat, a software platform that predicts intercellular communication networks based on expression data of known ligand-receptor pairs. Applying CellChat analysis to a lung scRNA-seq dataset from sham and AKI mice (Supplemental Figure 3) predicted neutrophil chemoattractant signals that may drive neutrophil train formation after AKI. The total number of predicted signaling interactions in the lung vascular niche between neutrophils, monocytes, transitional monocytes, interstitial macrophages, and endothelial cells modestly increased after AKI (∼5%) (Figure 6E). Cell-cell communication analysis comparing sham control with AKI predicted a strong relevance of classical monocyte to neutrophil signaling after AKI via the neutrophil chemoattractant CXCL2 acting on CXCR2 receptor expressed in neutrophils (Figure 6F). It also predicted that CXCL2 produced by neutrophils acts on neutrophils themselves in an autocrine loop. Injection of CXCL2-neutralizing antibody in AKI animals as compared to IgG control significantly suppressed neutrophil train formation in the lung (Figure 6G). These data suggest that CXCL2 released intravascularly by CCR2^+^ classical monocytes establishes a critical CXCL2 gradient within lung capillary vessels that is required to form neutrophil trains by attracting neutrophils to monocytes. This is further supported by our already presented intravital imaging findings at very early time points after AKI (Figure 2E).

### Comparative analysis of lung scRNA-seq data reveals that remote lung injury as compared to direct lung injury lacks alveolar macrophage activation and their release of neutrophil chemoattractants

To discern differences between direct and remote lung inflammation after AKI that may explain neutrophil extravasation and transmigration into alveoli in the former but not in the latter, we performed comparative scRNA-seq analysis of the transcriptomic landscape of lungs from control or syngeneic lung transplants, representing a model of direct lung injury (warm IRI) with high neutrophil extravasation and transmigration into alveoli (35), and of lungs from sham or AKI injured animals, notably lacking extravasation and transmigration into alveoli (Figure 1). We scaled, standardized, and integrated scRNA-seq data from mouse control lungs, syngeneic lung transplants 2 hours after transplant (36), together with our aforementioned lung scRNA-seq dataset from sham and AKI mice. After excluding doublets and cells with low-quality expression data, 73,814 lung cells underwent clustering using Seurat v4 (37).

When comparing scRNA-seq data of direct lung injury with that of AKI-induced remote lung inflammation relative to respective controls, we found that neutrophils in remote lung inflammation (exclusively intravascular) as compared to neutrophils in direct lung inflammation (mostly extravasated) show much higher innate immune signaling with upregulation of *Myd88*, a downstream effector of signaling via Toll-like receptors in response to damage-associated molecular patterns (DAMPs) (38), and of *IL1b*, a major effector of early Type 1 immune responses (39). In contrast, alveolar macrophages in direct, but not in remote lung inflammation showed significant activation of innate immune signaling, again with *Myd88* and *IL1b* upregulation, and strong expression of the neutrophil chemoattractant CXCL2 (Figure 7A). Consistent with this, analysis of differentially expressed genes (DEGs) showed that alveolar macrophages exhibit less leukocyte activation and cytokine signaling after AKI as compared to direct lung injury (Figure 7B). A comprehensive comparison of monocyte/macrophage populations in direct and remote lung inflammation also highlighted the lack of alveolar macrophage activation after AKI, while other cell types (interstitial macrophages, Ly6C^−^, and Ly6C^+^ monocytes) exhibited comparable or even more intense transcriptomic inflammatory signatures after AKI than after direct lung injury (Supplemental Figure 7). CXCL2 production by alveolar macrophages could establish a CXCL2 gradient directed towards the alveolus that may be responsible for the observed neutrophil extravasation and transmigration into alveoli in direct lung injury. Further, our analysis suggests that this alveolar macrophage derived extravasation signal is missing in AKI-induced remote lung inflammation. To assess whether neutrophils are in principle able to extravasate during AKI-induced remote lung inflammation, we instilled CXCL2 intratracheally just after sham or AKI. In comparison to vehicle control a one-time instillation of CXCL2 into the alveolus right after AKI significantly increased the rate of neutrophil extravasation by over 20-fold at 24 hours (0.41% to 11%) (Figure 7C). Alveolar CXCL2 was able to induce extravasation of comparably more intravascular neutrophils in sham mice as in AKI mice (Supplemental Figure 8), indicating that neutrophil stiffness, which is more prominent in AKI neutrophils (Figure 4), at least partially contributes to the neutrophil intravascular retention phenotype. These findings suggest that absence of alveolar macrophage activation and the lack of a neutrophil chemoattractant gradient into the alveolus likely contributes to the relative lack of neutrophil extravasation after AKI. A summary of our findings is shown in Figure 8.

**Figure 7.**
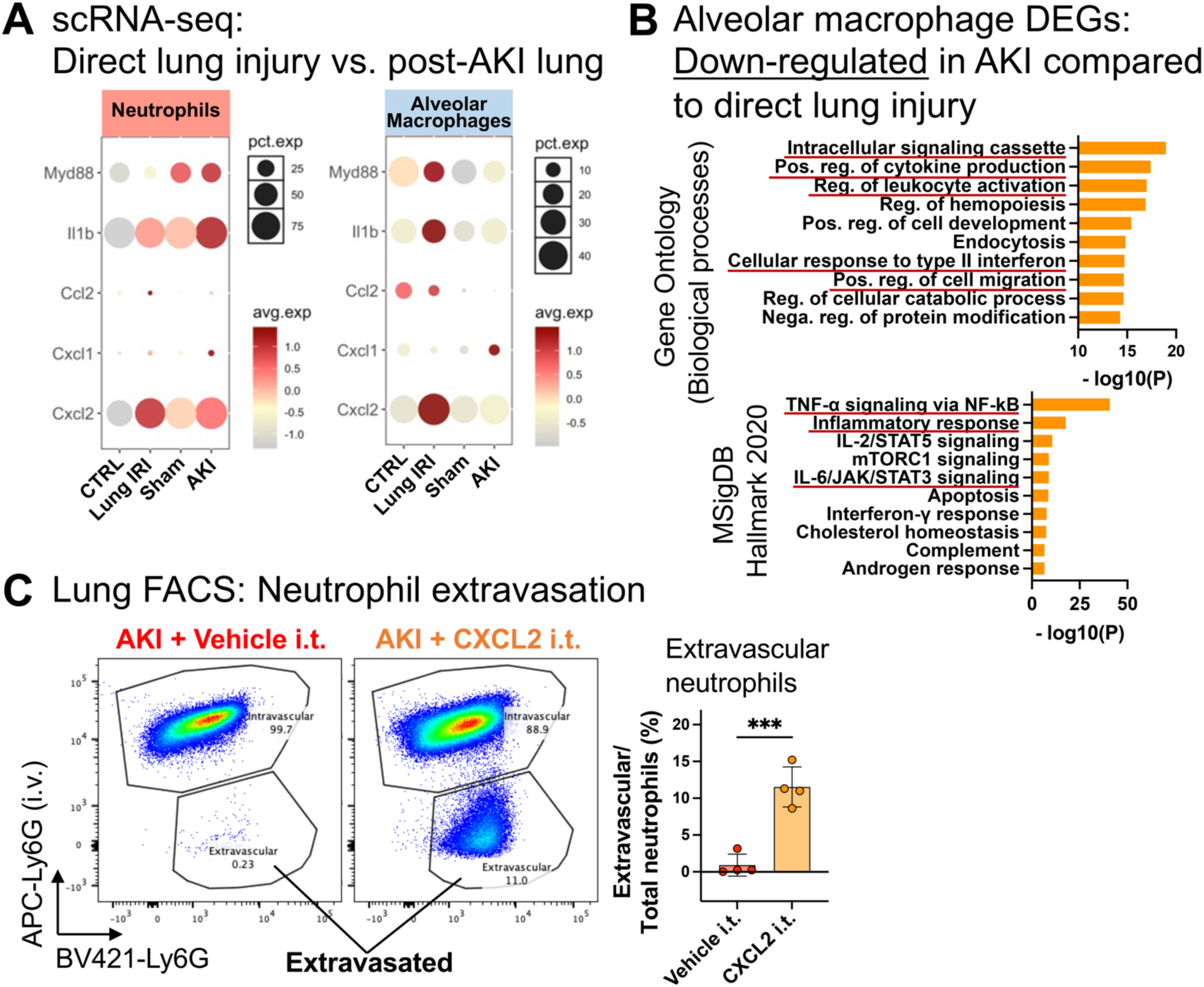
Comparative analysis of lung scRNA-seq data reveals that remote lung injury as compared to direct lung injury lacks alveolar macrophage activation and their release of neutrophil chemoattractants. (**A**) Comparison of inflammatory genes using single-cell RNA seq analysis: A lung syngeneic transplant dataset was analyzed as a model of direct lung injury and compared to our AKI dataset (both are warm IRI). Expression of inflammatory molecules in neutrophils and alveolar macrophages are shown. The dot size denotes percentage of cells expressing each gene, and the color scale represents average gene expression levels. (**B**) Differentially expressed gene (DEG) analysis in alveolar macrophages: downregulated genes in post-AKI lung compared to direct lung injury were analyzed using gene ontology (GO) and MSigDB Hallmark pathway analysis. GO terms and pathways related to inflammation and associated with activation of macrophages are underlined. (**C**) FACS quantification of extravasated neutrophils 24 hours after AKI. Either mouse CXCL2 protein (0.01 µg/g) or vehicle in 50 µL sterile saline was administered intratracheally immediately after AKI surgery. Data represent mean ± SD and n = 4 mice per group. ***P < 0.001

**Figure 8.**
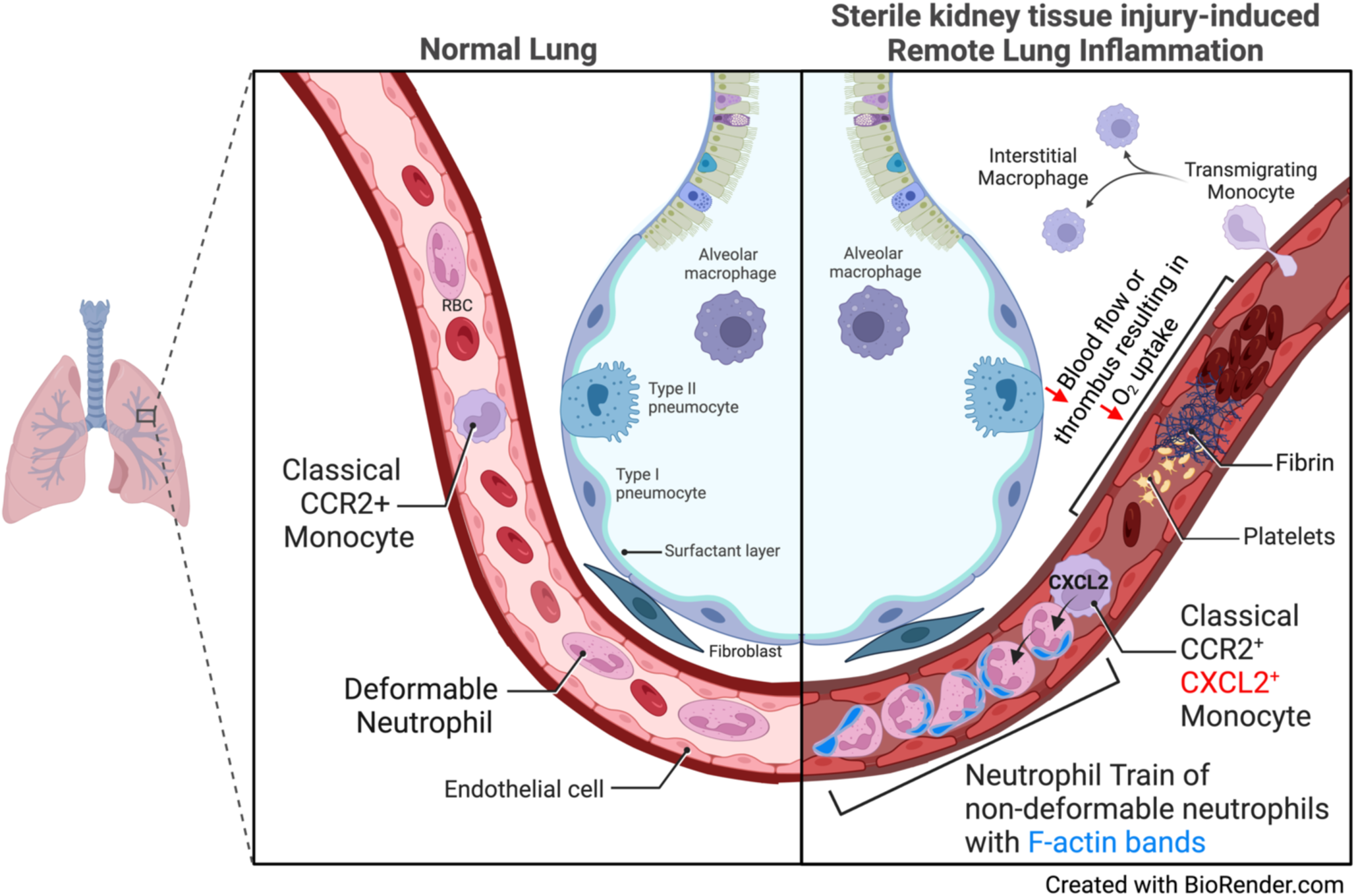
Schematic summary. AKI induces rapid intravascular “neutrophil train” formation in lung alveolar capillaries, a form of neutrophil retention. Rapid retention is enhanced by decreased deformability secondary to F-actin polymerization (submembrane F-actin bands) in circulating neutrophils that impedes their lung capillary passage. CCR2^+^ classical monocytes are required for neutrophil train formation and release CXCL2 to attract neutrophils into trains. Neutrophil train formation reduces alveolar capillary blood flow and is associated with thrombosis (thrombi contain both platelets and fibrin). This capillary perfusion defect leads to reduced oxygenation due to a ventilation perfusion mismatch, a scenario that differs from infectious inflammatory lung diseases, such as bacterial pneumonia or pulmonary alveolar edema where ventilation is affected.

## DISCUSSION

Our work reveals that lung neutrophil recruitment after sterile kidney tissue injury as compared to direct lung injury requires different cellular cues, localizes almost exclusively to the intravascular compartment, and is mechanistically linked to the development of hypoxemia by causing a lung capillary perfusion defect, rather than by affecting alveolar function and ventilation.

A role of monocytes and alveolar macrophages in lung neutrophil recruitment and their extravasation different from sterile AKI has been demonstrated in direct lung injury after bacterial challenge and during lung IRI, a process inherent to pulmonary transplantation (35). Here, intravascular lung non-classical monocytes are activated by lung tissue injury-released DAMPs and drive neutrophil accumulation in lung vessels by production of the neutrophil chemoattractants CXCL1 and CXCL2 (19, 29). At the same time non-classical monocytes produce IL1β which activates tissue resident alveolar macrophages to produce the monocyte chemoattractant CCL2. CCL2 recruits CCR2^+^ classical monocytes into lung vessels, a requirement for the observed massive neutrophil extravasation into lung tissue and alveoli during direct injury (30) (reviewed in (40)). In contrast, our results show that non-classical monocytes and alveolar macrophages are dispensable for AKI-induced remote lung neutrophil accumulation. Alveolar macrophages did not show activation of innate immune signaling pathways during AKI, as opposed to direct lung injury, suggesting that AKI-induced innate immune activating signals from the circulation are unable or not strong enough to reach alveolar macrophages. However, alveolar macrophages do play a role in sepsis-induced remote lung inflammation, another form of indirect lung injury. In sepsis from extrapulmonary infections alveolar macrophages become activated and drive significant neutrophil extravasation into the alveolar space (41). Such alveolar neutrophil extravasation is in fact also observed in septic patients with ARDS (septic ARDS) (42) (reviewed in (43)), suggesting that sepsis induces additional signals that reach and activate alveolar macrophages or that during sepsis intravascular inflammatory and innate immune activating signals are more pronounced than during AKI. Mimicking a neutrophil chemoattractant signal emanating from the alveolus, intratracheal instillation of the neutrophil chemoattractant CXCL2 induced extravasation of neutrophils after AKI, and even better after sham. This shows that non-deformability of neutrophils does not principally interfere with extravasation, but that better deformability as seen in sham neutrophils enhances extravasation in the presence of an alveolar chemoattractant signal. Thus, neutrophils have the capacity to extravasate after AKI, but lack the alveolar signals to do so. CCR2^+^ monocytes are drivers of neutrophil extravasation into injured tissues not only in direct lung injury (31), but also for example during arthritis, as shown with intravital imaging (34). We can only speculate why CCR2^+^ monocytes do not exert this function in remote lung inflammation. Perhaps alveolar macrophage activation is necessary in addition to CCR2^+^ monocyte activation to mediate neutrophil extravasation, as is observed in direct lung injury.

We and others have previously reported on the lack of neutrophil transmigration into alveoli up to 24 hours after AKI based on bronchoalveolar lavage analysis in mice (6, 44). This suggested that neutrophils may not significantly extravasate and/or transmigrate through the pulmonary epithelial barrier during remote lung inflammation, in mice or humans. Enhanced neutrophil accumulation in lung vessels 24 hours after AKI in mice has been observed, but lung tissue validation was limited, and its physiological significance was not studied (9). The normal lung circulation is unique in that it contains marginated intravascular neutrophils and also classical and non-classical monocytes, together forming an immune niche poised to rapidly respond to various immune challenges, in particular invasion of pathogens (45). The entire cardiac output passes through the lungs, such that during dissemination of infection via the bloodstream pathogens have to pass through the lungs. Like sepsis, AKI induces a systemic inflammatory response syndrome (SIRS) involving many of the same circulating pro-inflammatory mediators and also an increase of circulating neutrophils and monocytes (7). Pathogens in the bloodstream induce the formation of sizable intravascular neutrophil swarms in larger caliber lung vessels that facilitate pathogen capture and clearance and serve as an important evolutionarily conserved defense mechanism against sepsis (46). Using intravital lung imaging, we show that AKI induces a different form of intravascular neutrophil accumulation, namely intravascular neutrophil retention in “neutrophil trains”. Lung neutrophil trains impede blood flow in alveolar capillaries, sites where an intact circulation is required for oxygen exchange. Our work reveals that capillary neutrophil trains are able to reduce capillary blood flow to a degree that rapidly affects oxygenation after AKI (within 2–6 hours), even without significant interstitial or alveolar edema and alveolar exudates. Based on our previous observation that lung neutrophil accumulation reaches a plateau at 12–24 hours after AKI (6), neutrophil train formation and lung capillary flow impediment likely progress significantly after the 2 hours intravital imaging time point we obtained (Figure 2A) but becomes critical for oxygenation somewhere between 2–6 hours after AKI. A possible correlate of reduced capillary blood flow, AKI-induced red blood cell “rouleaux” formation in lung vessels has been reported previously, but its effects on blood flow or physiological outcomes was not determined (47). We speculate that lung capillary intravascular neutrophil train formation and slowed lung capillary blood flow after sterile kidney tissue injury represent an adaptive reaction that may serve to sequester activated immune cells and circulating cytokines in the lung, protecting the organism from their overt systemic action, but at the expense of causing blood oxygenation deficits that may result in hypoxemia (maladaptive consequence). Extensive sepsis-induced neutrophil swarms can indeed also negatively impact oxygenation (48). Widespread medium to large vessel arterial and venous thrombosis containing neutrophil macroaggregates has been described after gut ischemia in mice, and also in patients with severe ARDS (49).

We show that rapid lung neutrophil capture after AKI is enhanced by an induced reduction in neutrophil deformability. Changes in neutrophil deformability (23) and lung retention lasting >40min has been observed in humans following the injection of *ex vivo* primed neutrophils (50), and changes in flow and adhesive properties (“stiffening”) of circulating immune cells have also been documented in humans with trauma (24), ARDS (51), cardiac tissue injury (52), or sepsis (53, 54). Canonical steps of neutrophil recruitment that mediate crawling and arrest, such as LFA-1/ICAM1 or Mac-1/ICAM1 neutrophil/endothelial interactions, are not required in remote lung inflammation. This is not unprecedented as neutrophil recruitment is strongly context dependent (46) and does not always involve integrin activation, and non-canonical pathways involved in neutrophil recruitment have been described in different tissues (55).

In several contexts, leukotriene-B-4 (LTB-4) is required for the initiation of intravascular neutrophil swarming and clustering behavior (reviewed in (46)), in particular in fungal sepsis (48). However, we did not find any significant expression of LTB-4 receptors in lung cells after sterile kidney tissue injury. Instead, our model supports the involvement of the neutrophil chemoattractant CXCL2 released by classical monocytes as the signal that assembles captured neutrophils into “trains” in the lung early after AKI. At later time points, as our lung cell-cell communication analysis on day 1 after AKI suggests, neutrophils produce CXCL2 themselves, likely contributing to increased “train” length and their retention in the lung. CXCL2 has not been directly linked to lung neutrophil recruitment after AKI, but mice globally deficient in the CXCL1/2 receptor CXCR2 show reduced remote lung neutrophil recruitment after AKI (56), supporting our data. Other source cells and lung neutrophil chemoattractants in AKI-induced remote lung inflammation have been identified. For example, neutrophil accumulation has also been linked to AKI-induced CXCL1 produced by lung endothelial cells. Inhibition of IL-6 in *wt* mice reduced remote lung inflammation, lung CXCL1 expression, and lung neutrophil accumulation after AKI (10). IL-6 injection into IL-6-deficient mice with AKI induced lung CXCL1 expression and lung neutrophil accumulation; CXCL1-neutralizing antibody or CXCR2 global knockout conferred protection. Lung CXCL1 protein colocalized with endothelial cell markers in lung tissue, suggesting that CXCL1 may be produced by endothelial cells after AKI (56). Lung neutrophil recruitment driven by endothelial CXCL1 may act in concert with the CXCL2 mechanism described in this manuscript. However, our scRNA-seq analysis 24 hours after AKI compared to sham controls did not reveal significant CXCL1/2 expression in lung endothelial cells (6).

While lung edema can occur after AKI due to volume overload with left ventricular failure in the setting of oliguria (cardiogenic edema), AKI patients with respiratory compromise frequently do not exhibit signs of volume overload such as elevated pulmonary artery wedge pressure on cardiac catheterization (57, 58), and also do not significantly improve after volume removal using dialysis (12, 59). Instead, non-cardiogenic lung edema driven by endothelial injury in response to AKI-released circulating inflammatory factors resulting in endothelial leak with interstitial and alveolar edema has been proposed as the dominant mechanism of AKI-induced hypoxemia. Interstitial wall thickness increases, and alveolar edema would interfere with ventilation and oxygenation by increasing the distance between the ventilated space and red blood cells traveling in capillaries (reviewed in (4, 5, 16)). Our findings provide an alternative or at least additional explanation for respiratory failure in the context of AKI that does not improve with fluid removal, whether AKI occurs in isolation or as part of existing respiratory failure from primary or secondary causes (such as COVID-19, bacterial pneumonia, sepsis or trauma). The lack of significant interstitial/alveolar edema and of inflammatory alveolar exudates after AKI likely explain why such patients frequently show only subtle changes on their chest radiographs that cannot explain the observed degree of hypoxemia. Strategies to detect and prevent remote lung inflammation and capillary neutrophil train formation may represent novel therapeutic avenues to diminish or prevent AKI-induced respiratory complications in the clinic. Since the observed lung effects develop rapidly and AKI is often detected late in the clinic, efforts to understand how neutrophil trains can be detected, de-primed, dissolved and released from the lung appear particularly important points to address in future research in critically ill patients with AKI.

## METHODS

### Sex as a biological variable

Our study used male C57BL/6 mice because female mice exhibit high resistance against ischemic injury, show high mortality, and large variability in our measurements. However, we have also observed the kidney-lung phenotype we describe here in female mice. We thus expect our findings in male mice presented here to be relevant for both sexes.

### Animal experiments

8–12-week-old male C57BL/6 (strain #000664) and NR4A1-KO (#006187) mice were purchased from The Jackson Laboratory (Bar Harbor, ME) and kept in 12 hours-light/dark cycle with liberal water and standard chow diet. In our AKI model 20-minutes of bilateral ischemia-reperfusion (IR) injury was introduced at 37°C using the flank approached as previously described (6). Sham operation was performed using the same surgical procedure without clamping the renal hilum with surgical clips. In select experiments, following intubation with a 20G catheter, 0.01 µg/g of lipopolysaccharide (LPS) derived from *Pseudomonas aeruginosa* (Sigma-Aldrich, #L8643), or 0.01 µg/g of mouse CXCL2 (R&D, #452-M2-010), dissolved in 50 µL of sterile saline was intratracheally instilled. For the neutrophil depletion experiment, mice were treated with 400 µg of anti-Ly6G antibody (BioXCell, #BE0075) or the same amount of rat IgG isotype control 24 hours before and immediately before AKI surgery. For CCR2^+^ monocyte depletion, C57BL/6 mice were injected with anti-CCR2 antibody (MS-21) 20 µg per day for four consecutive days (33, 60) prior to IR injury. For CD18 and CXCL2 inhibition, 200 µg of anti-CD18 (BioXCell, #BE0009) and 100 µg of anti-CXCL2 monoclonal antibody (Invitrogen, #MA5-23737) was injected, respectively (half the dose i.p. 1 hour prior to ischemia and the other half i.v. at reperfusion). Control animals were treated with the same dosage of rat IgG isotype (Invitrogen, #10700). The organs were harvested 2–24 hours after IR injury, as specified in each experiment. In the blood gas measurement experiment, mice were intubated following injections of ketamine (90 µg/g) and xylazine (10 µg/g), then mechanically ventilated with 100% O_2_. Arterial blood samples were directly collected from the ascending aorta using a syringe with 30G needle under a surgical microscope. These samples were analyzed within 1 minute using the i-STAT portable blood gas analyzer system (Abbott, IL).

### Single-cell preparation from mouse lung

Twenty-four hours after sham or AKI surgery, mice were euthanized and perfused with 20 ml of ice-cold PBS from the left and right ventricles to remove remaining blood in the lung vasculatures. Perfused lung was immediately dissected and chopped into <1mm^3^ dices by scalpels and razors. For single cell preparation samples were incubated for 60-min at 37°C with gentle shaking in RPMI media supplemented with 50 mg/ml of Liberase TL (Roche, #5401020001), 40 U/ml of DNAse I (Sigma-Aldrich, #D4527), and 0.75 mg/ml of hyaluronidase (Sigma-Aldrich, #H3506). At the end of incubation each sample was filtered twice, once through a 70 mm and once through a 40 mm strainer. Red blood cells in the samples were removed by ACK lysing buffer (Quality Biological, #118156101). Single-cell suspensions were then washed and resuspended in FACS buffer (PBS, 0.5% BSA, 2mM EDTA, 0.01% NaN_3_), and used for single-cell RNA sequencing (scRNA-seq) and/or flow cytometry analysis.

### Single-cell RNA sequencing (scRNA-seq) and data analysis

scRNA-seq analysis was conducted on lung and kidney samples from sham and AKI as previously described (6). Briefly, a total of six samples were pooled for each group and live cells were sorted using propidium iodide staining using a FACSAria III (BD Biosciences) flow cytometer. cDNA libraries were prepared using the Chromium Single Cell 5’ Library Kit v2 (10x Genomics, Pleasanton, CA) following manufacturer’s instructions. The amplified full-length cDNA libraries were then submitted to the Genome Technology Access Center of Washington University for sequencing, with a sequencing depth of 50,000 reads. All subsequent data processing steps were performed using R software (v4.1.0) with Seurat v4 (37). Expression matrices for each sample were imported into R as Seurat objects. Quality control was initially performed by retaining cells with gene expression counts ranging from 200 to 3200. Cells with more than 10% mitochondrial genes and genes not expressed in at least three cells were excluded. After these quality control steps, the SCTransform and Harmony functions were employed for normalization, scaling, variance stabilization, and integration of the datasets (61, 62). After quality control and integration, we performed clustering by applying a K-nearest neighbor graph with a resolution of 0.2 on the result of principal component analysis (PCA). We visualized the clustering using uniform manifold approximation and projection (UMAP). To assign cluster identities, we first compiled a list of lung and kidney cell types and their currently established markers (63, 64) and performed manual annotation using those markers and additional differentially expressed genes (DEGs) between clusters identified by the FindAllMarkers() function of Seurat. The expression levels of genes of interest in each cluster were visualized using ‘plot1cell’ package (65). Cell-cell communication analysis within the scRNA-seq datasets was performed using CellChat (66). Functional properties of statistically significant DEGs (P < 0.05) were then investigated by GO term analysis using Metascape (67) and MSigDB Hallmark 2020 Pathway analysis using Enrichr (68).

### Flowcytometry analysis

Single-cell suspensions were stained with LIVE/DEAD Aqua cell dye (Invitrogen, #L34965, dilution 1:200) for 30 minutes. After washing with PBS, FcR blocking with anti-mouse-CD16/32 antibody (BioLegend, #101302, dilution 1:100) was performed for 15 minutes in FACS buffer, followed by 30-min incubation with the following diluted antibodies: BV605-anti-mouse CD45 (BioLegend, #103139, 1:200), BV421-anti-mouse Ly6G (BioLegend, #127627, 1:200), BUV395-anti-mouse CD11b (BD Biosciences, #563553, 1:200), BV786-anti-mouse SiglecF (BD Biosciences, #740956, 1:200), FITC-anti-mouse F4/80 (Invitrogen, #11480181, 1:100), PE-anti-mouse MERTK (BioLegend, #151506, 1:100), APC-anti-mouse Ly6C (BioLegend, #128016, 1:400). Unstained and single stained compensation samples were prepared from live lung or spleen cells for each FACS run. For quantification of F-actin in neutrophils, blood leukocytes were incubated with rhodamine-conjugated phalloidin (Invitrogen, #R415, 1:200) for 30 minutes at room temperature after permeabilization by 0.2% Tween-20. FACS analysis was performed on BD Fortessa X20 with five lasers and data was analyzed with FlowJo software (BD, NJ).

### In vivo neutrophil extravasation assay

In order to distinguish intravascular from extravasated neutrophils, 10 µL of APC-anti-mouse-Ly6G antibody (BioLegend, #127614) was intravenously injected 10 minutes before sacrifice as previously reported (17). After dissociation of lungs into single cell suspensions, each sample was stained by another anti mouse Ly6G antibody (BioLegend, #127627) carrying a different fluorescence (BV421). During FACS analysis, intravascular neutrophils were identified as double positive (APC^+^, BV421^+^), whereas extravasated neutrophils were single positive (APC^+^, BV421^−^) since they were not reached by the intravenously applied first antibody.

### Intravital imaging of the lung

Intravital lung imaging in CCR2-GFP mice was performed using two-photon microscopy as previously described (35). Briefly, 1.5 hours after IR or sham surgery, mice were orotracheally intubated with a 20 G catheter and ventilated with 1.5% isoflurane in air supplemented with 2L/min of oxygen (tidal volume: 220–250 µl, respiratory rate 250/min). To visualize neutrophils after AKI or sham, blood vessels, and blood flow, 8 µL of PE-anti-mouse-Ly6G antibody (BioLegend, #127608), 10 µL of 655-nm Q-dots (Invitrogen, #Q21021MP), and 5 µL of 1-µm fluorescent beads (Invitrogen, #F13080) were intravenously injected prior to imaging. A thoracotomy with partial removal of 4–5 left lower ribs exposed enough lung tissue for two-photon microscope imaging. Our strategy allowed intravital lung imaging at about 2 hours after sham or AKI. The video rate was set to 300 msec and six random areas of the lung were recorded for at least 25 seconds per animal. The entire surgical and acquisition process was conducted on a heated pad to maintain the body temperature of mice at 37°C. To obtain intravital images of a comparable ischemia-reperfusion-induced direct lung injury model, we used lung tissue with a warm ischemia time of 45 min at 28°C, transplanted into syngeneic LysM-GFP mice. Intravital imaging was performed at the same imaging facility 2 hours after transplantation; detected GFP-positive cells are almost exclusively monocytes (19).The acquired videos were analyzed using the Imaris software version 9.8.2 (Oxford Instruments, Abingdon, UK).

### Histology and immunofluorescence staining

For histological analysis, lung samples were placed inside a 20-mL syringe and inflated by creating negative pressure with pulling the syringe plunger, repeated several times until the lobes sink in 4% paraformaldehyde. After inflation, the lobes were fixed in 4% paraformaldehyde overnight at 4°C, then processed and embedded in paraffin. Alveolar wall thickness was examined with 4-µm sections from paraffin embedded lung samples using standard hematoxylin and eosin (H&E) staining kit (Sigma, #HT107/HT109) as previously reported (6, 69). In short, alveolar walls in randomly selected lung regions that met the grid set by ImageJ were manually measured for wall thickness, and the average of at least 100 measurements was recorded as an observation from each sample. Immunofluorescence staining was performed on 7-µm fresh-frozen sections embedded into OCT compound (Fisher, #4585). In select experiments, 25 µL of 1-µm fluorescent microbeads (Invitrogen, #F13080) were intravenously injected 10 minutes prior to sacrifice to assess stagnation in the microcirculation in the lung. Six images/lung were collected for blinded quantification by experienced lab members. OCT tissue cryo-sections were washed with PBST (0.05% Tween-20 in PBS) for 5 minutes at room temperature and permeabilized with 0.1% Triton-X in PBS for 5 minutes, followed by PBST washes (3 x 5 min). After 30 min incubation in blocking buffer (1% BSA and 10% normal goat serum in PBS), sections were incubated with primary antibodies overnight at 4°C, followed by PBST washes (3 x 5 min), 1 h incubation at room temperature with secondary antibodies (where appropriate), 5 min with Hoechst dye (Thermo, #62249), followed by PBST washes (3 x 5 min) and mounting solution (2 mg/mL para-phenylenediamine) was applied. Primary Abs: anti-mouse-Gr-1 (FITC*-*conjugated, eBioscience, #14-5931), rabbit anti-mouse-CD68 (Abcam, #ab125212), rat anti-mouse-CD41a (Invitrogen, #14-0411-82), and rabbit anti-mouse-Fibrinogen (Abcam, #ab34269). Secondary Ab: Alexa Fluor 488 donkey anti-rat-IgG (Invitrogen, #A21208), Alexa Fluor 594 goat anti-rabbit-IgG (Thermo, #A11037), Alexa Fluor 546 donkey anti-rabbit-IgG (Invitrogen, #A10040). For quantitative analysis, we selected at least five representative areas that were captured with a Nikon Eclipse E800 fluorescence microscope at a 200x magnification. In select experiments, F-actin in neutrophils was stained by rhodamine-conjugated phalloidin (Invitrogen, #R415) for 30 minutes at room temperature following fixation and permeabilization using 0.2% Tween-20.

### Statistics

All results are reported as the mean ± SD. Comparison of 2 groups was performed using an unpaired, 2-tailed t-test. Comparison of 3 or more groups was performed by one-way ANOVA, followed by Tukey’s post hoc test. Statistical analyses were performed using GraphPad Prism 10.1.1 for macOS (GraphPad Software, MA). *P* values less than 0.05 were considered significant.

### Study approval

All animal experiments were approved by Institutional Animal Care and Use Committee of Washington University School of Medicine (protocol #19-0044 and #22-0105).

## Data availability

The mouse scRNA-seq datasets used in this study have been uploaded and are available under the GEO accession number GSE249928 (AKI-lung) and GSE249242 (lung transplant). All data presented in the graphs are provided in the Supporting Data Values file in the Supplemental material.

## Authors contribution

AH conceived and coordinated the study. YK designed, performed, and analyzed most of the experiments. LN, CL, and AS performed part of the animal experiments, tissue sectioning and staining. YK, WL, and MM performed the intravital imaging. SK performed arterial blood collection under ventilation from the mice. HS and DK provided the scRNA-seq dataset of murine lung transplantation. AH and YK interpreted all the data, prepared the figures, and wrote the manuscript. EK, MM, and DK critically revised it. AH and MM provided experimental resources. AH supervised the entire project. All authors read and approved the final version of the manuscript.

## Acknowledgements

Anti-CCR2 antibody used for the in vivo monocyte depletion experiment was a generous gift from Prof. Dr. Matthias Mack (Universität Regensburg, Germany). The CCR2-GFP mice and CD169-DTR mice were provided by Prof. Drs. Steven Brody and Kory Lavine, respectively (both at Washington University in St. Louis). We appreciate Prof. Drs. Kory Lavine and Regina Clemens (Washington University in St. Louis) for useful advice from their expertise. We acknowledge the In Vivo Imaging Core (IVIC), at the Washington University School of Medicine, for two-photon imaging and multi-dimensional data analysis. YK was supported by the Japan Society for the Promotion of Science (JSPS) Overseas Research Fellowship (#202360162) and The Uehara Memorial Foundation Overseas Research Fellowship (#202140023). EK was supported by the American Heart Association (Career Development Award 20CDA35320006) and the American Society of Nephrology (KidneyCure Carl W. Gottschalk Research Scholar Grant). MM was supported by National Institute of Allergy and Infectious Diseases (R01AI077600). AH was supported by National Institute of Diabetes and Digestive and Kidney diseases (R01DK121200) and VA merit award (I01BX005322).

