## Supplemental Figures 1-8 for "Acute kidney injury triggers hypoxemia by inducing intravascular neutrophil retention that reduces lung capillary blood flow"

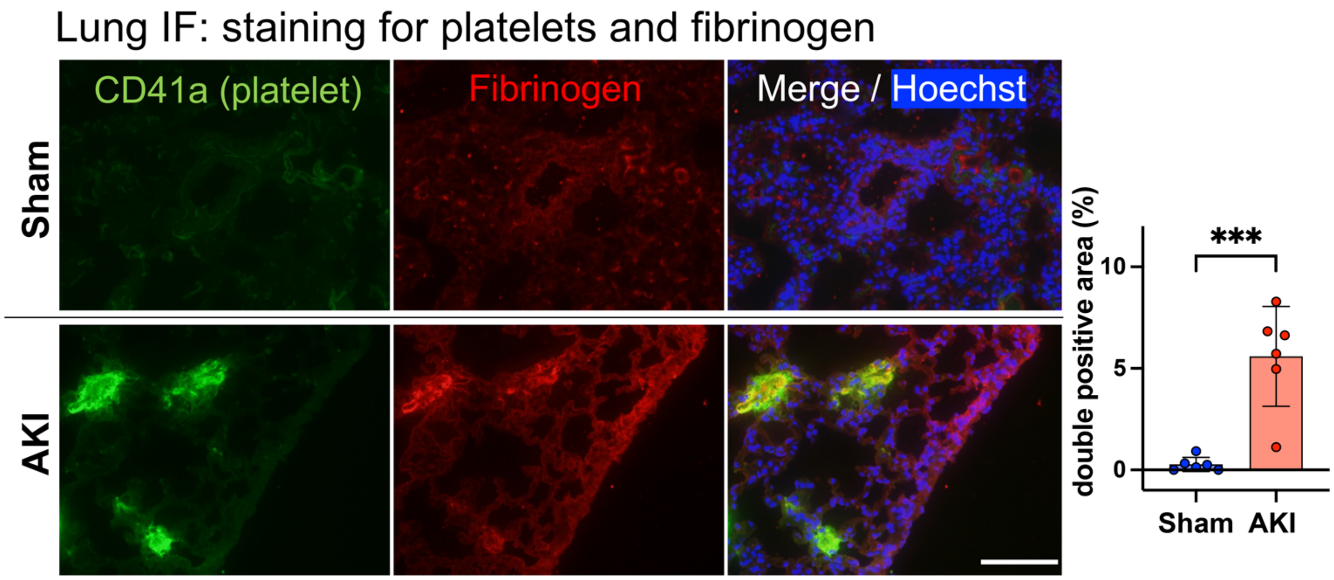


**Supplemental Figure 1. Capillary thrombosis in the lung after AKI.** Immunostaining for CD41a$^{+}$ lung platelets (green) and fibrinogen (red); graph shows quantification of double positive area after sham or AKI. Hoechst 33342 dye (blue) was used to visualize nuclei. Scale bar = 100 µm. Data represent mean ± SD and n = 6 mice per group. ***P < 0.001 compared to sham.

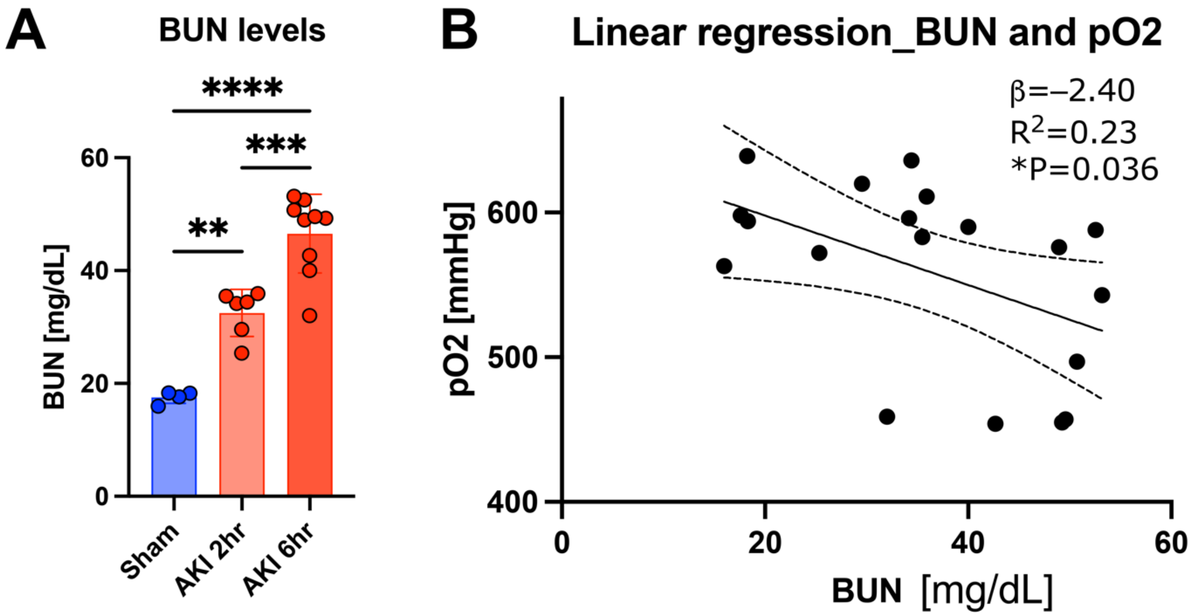


**Supplemental Figure 2. Association between blood urea nitrogen (BUN) and oxygenation.** (**A**) Serum BUN values after sham or 2–6 hours after AKI. (**B**) Linear regression between arterial pO2 [mmHg] and BUN. Arterial blood samples were collected from ascending aorta under surgical microscope following intubation and mechanical ventilation with 100% $FiO_2$. Total n = 19, *P < 0.05.

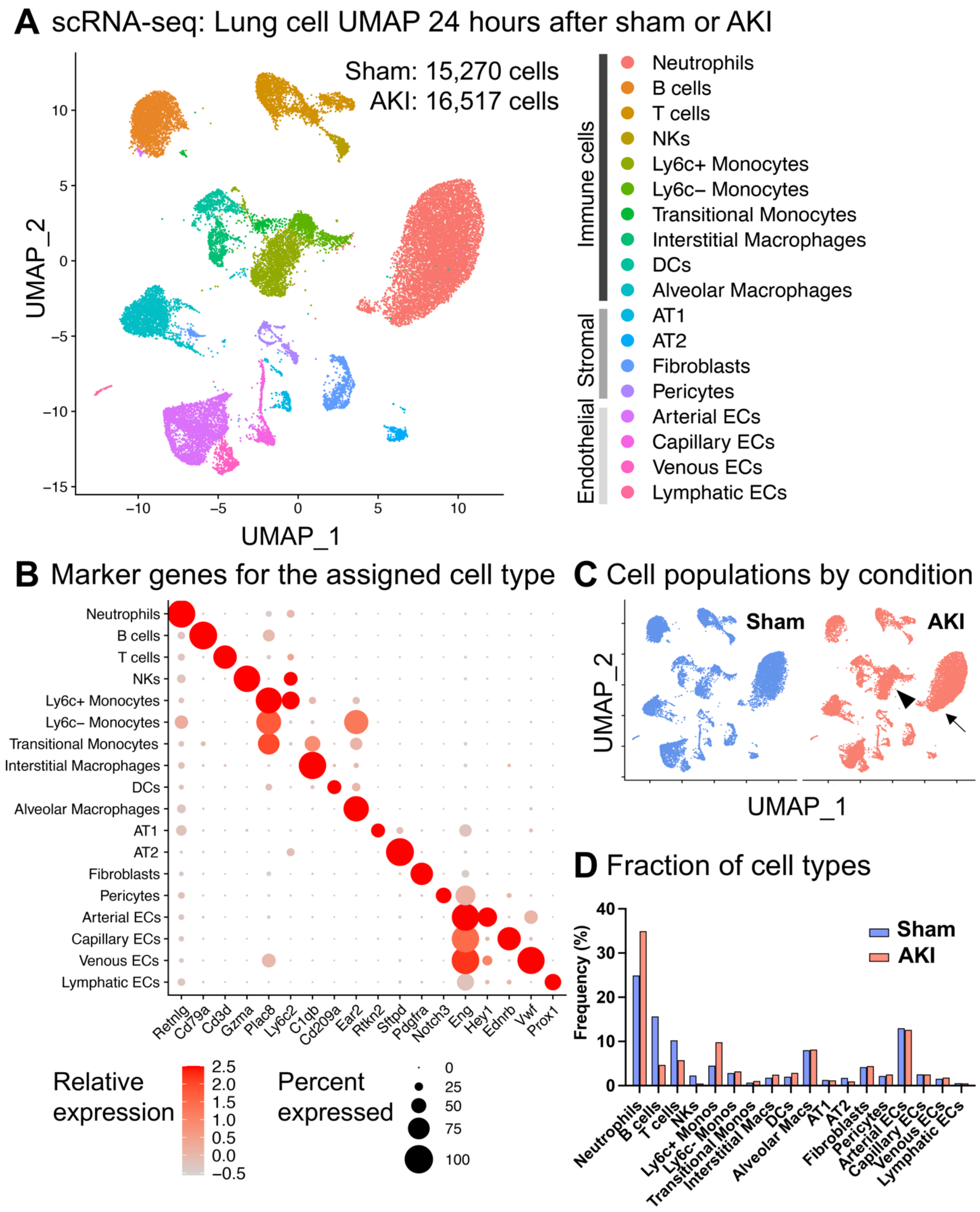


**Supplemental Figure 3. Single-cell transcriptomic landscape of remote lung inflammation induced by ischemic acute kidney injury.** (**A**) As an entry point to better understand cellular events involved in remote lung inflammation after AKI, we analyzed the transcriptomic landscape of lungs after ischemic acute kidney injury (AKI) at single-cell resolution. We scaled, standardized, and integrated single-cell

RNA sequencing (scRNA-seq) data from mouse lungs 24 hours after sham surgery or after bilateral renal ischemia-reperfusion injury-induced AKI. After excluding doublets and cells with low-quality expression data, 31,787 lung cells underwent clustering using Seurat v4 (1). We identified 18 lung cell types using canonical marker genes as previously reported (2): 10 different immune cell types, alveolar epithelial cell type 1+2 (AT1+2), fibroblasts, pericytes, and 4 endothelial cell types (arterial, capillary, venous, lymphatic). (**B**) Cell types and marker genes used for their identification were as follows: Neutrophils (*Retnlg*), B-cells (*CD79a*), T-cells (*Cd3d*), natural killer cells (*Gzma*), classical monocytes (*Plac8*$^{+}$, *Ly6c2* high), non-classical monocytes (*Plac8*$^{+}$, *Ly6c2* low), transitional monocytes (expressing both monocytes (*Plac8*) and interstitial macrophages (*C1qb*) markers), dendritic cells (*Cd209a*), alveolar macrophages (*Ear2*), AT1 (*Rtkn2*), AT2 cells (*Sftpd*), and fibroblasts (*Pdgfra*). Three separate endothelial cell clusters with high expression of *Eng* could be assigned to three different regions of the lung vasculature using *Hey1*, *Ednrb*, and *Vwf* as markers for arterial, capillary, and venous endothelial cells, respectively (3). Lymphatic endothelial cells were identified by high *Prox1* expression. (**C**) The lung cell types observed after AKI largely overlapped with those in the sham group; however, we identified several significant differences within the immune cell compartment, in particular in the neutrophil (small arrow) and monocyte/macrophage cluster (large arrowhead). (**D**) The overall frequency of neutrophils and Ly6c$^{+}$ classical monocytes increased in the lung following AKI, with a trend of increase in transitional monocytes and interstitial macrophages, whereas the proportion of lymphocytes decreased compared to the sham condition.

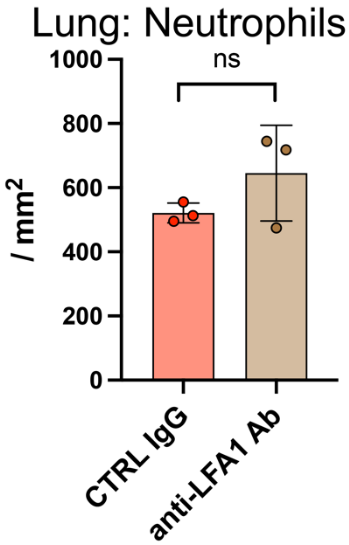


**Supplemental Figure 4. Anti-LFA1 neutralizing antibody did not prevent neutrophil accumulation in the lung after AKI.** Anti-LFA-1 antibody blockade vs IgG isotype control in C57BL/6 mice: A total of 100 µg of antibodies (anti-CD11a: BioXCell, #BE0006; rat IgG: Invitrogen, #10700) was injected (50 µg i.p. 1 hour prior to ischemia and another 50 µg i.v. at reperfusion). Quantification of lung neutrophils by Ly6G immunofluorescence staining after AKI. N = 3 mice/group. Data represent mean ± SD. Ns: not significant.

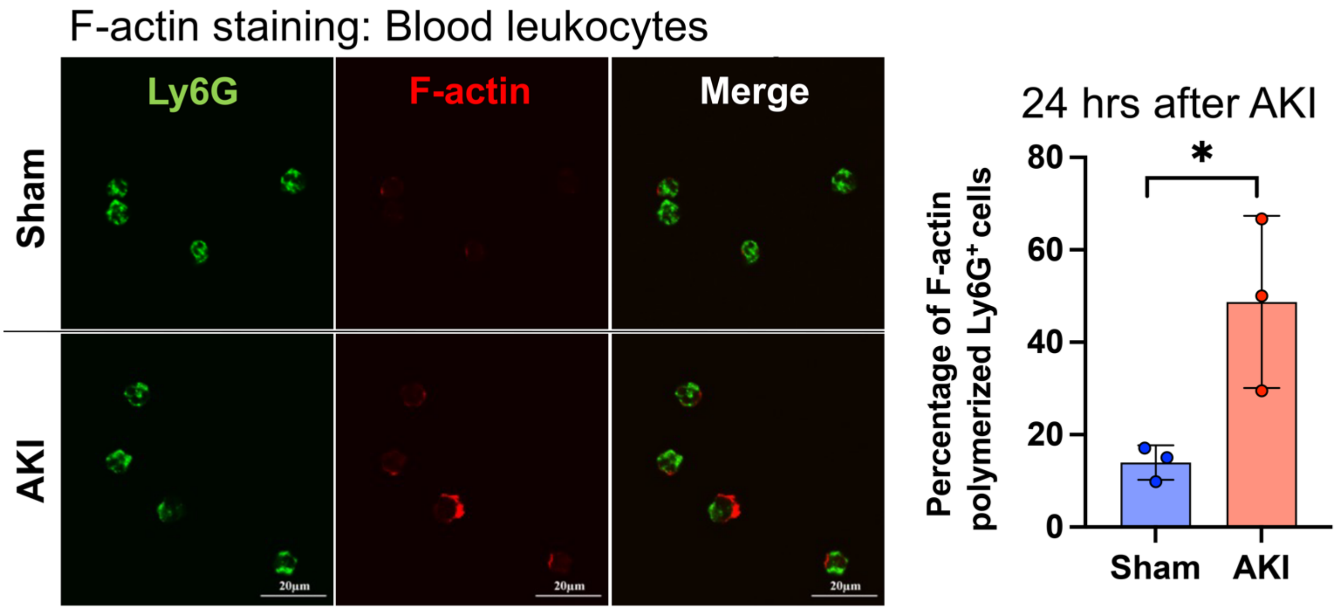


**Supplemental Figure 5. F-actin staining of the circulating neutrophils 24 hours after sham or AKI.** Neutrophils isolated from blood of sham or AKI animals (C57BL/6 mice) were stained with fluorescently-labeled phalloidin. Ly6G staining was used to distinguish neutrophils from other blood cell types. Proportion of F-actin$^{+}$ neutrophils is shown in the graph. N = 3 mice/group. Data represent mean ± SD. *$P < 0.05$.

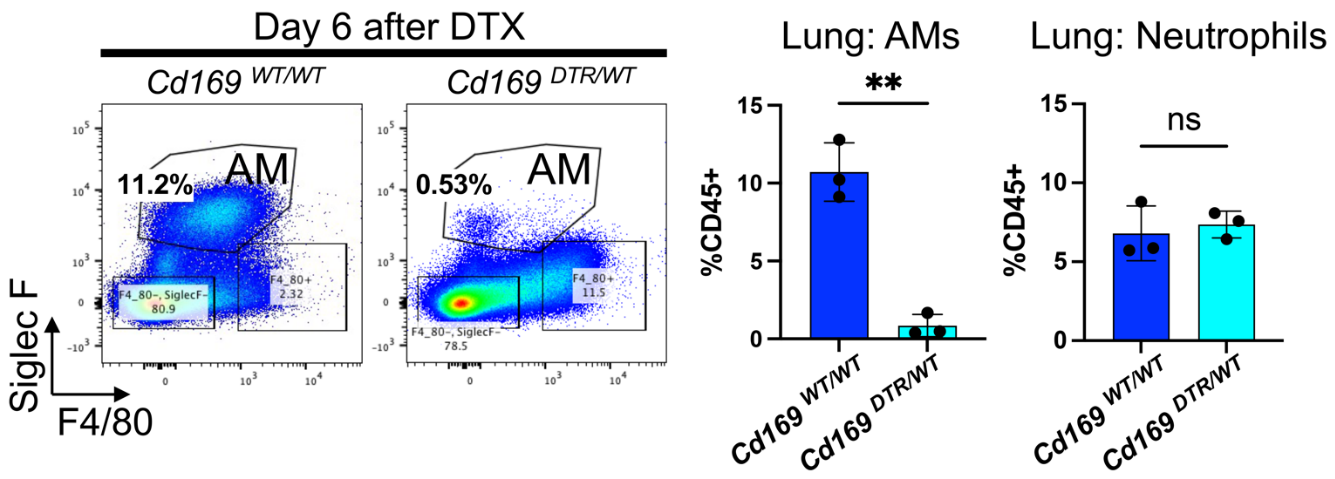


**Supplemental Figure 6. Baseline assessment of alveolar macrophage (AM) depletion model: Diphtheria toxin to CD169-DTR mice.** CD169-DTR heterozygous (*CD169*$^{DTR/WT}$) mice were analyzed alongside their wildtype (*CD169*$^{WT/WT}$) control littermates. Six days after intratracheal instillation of diphtheria toxin (DTX), lung single-cell suspensions were analyzed using FACS for baseline assessment (also see the schematic in Figure 5D). Lung alveolar macrophages were sufficiently depleted (>90% depletion), while the number of lung neutrophils remained unchanged. n = 3 mice/group. Data represent mean ± SD. **P < 0.01.

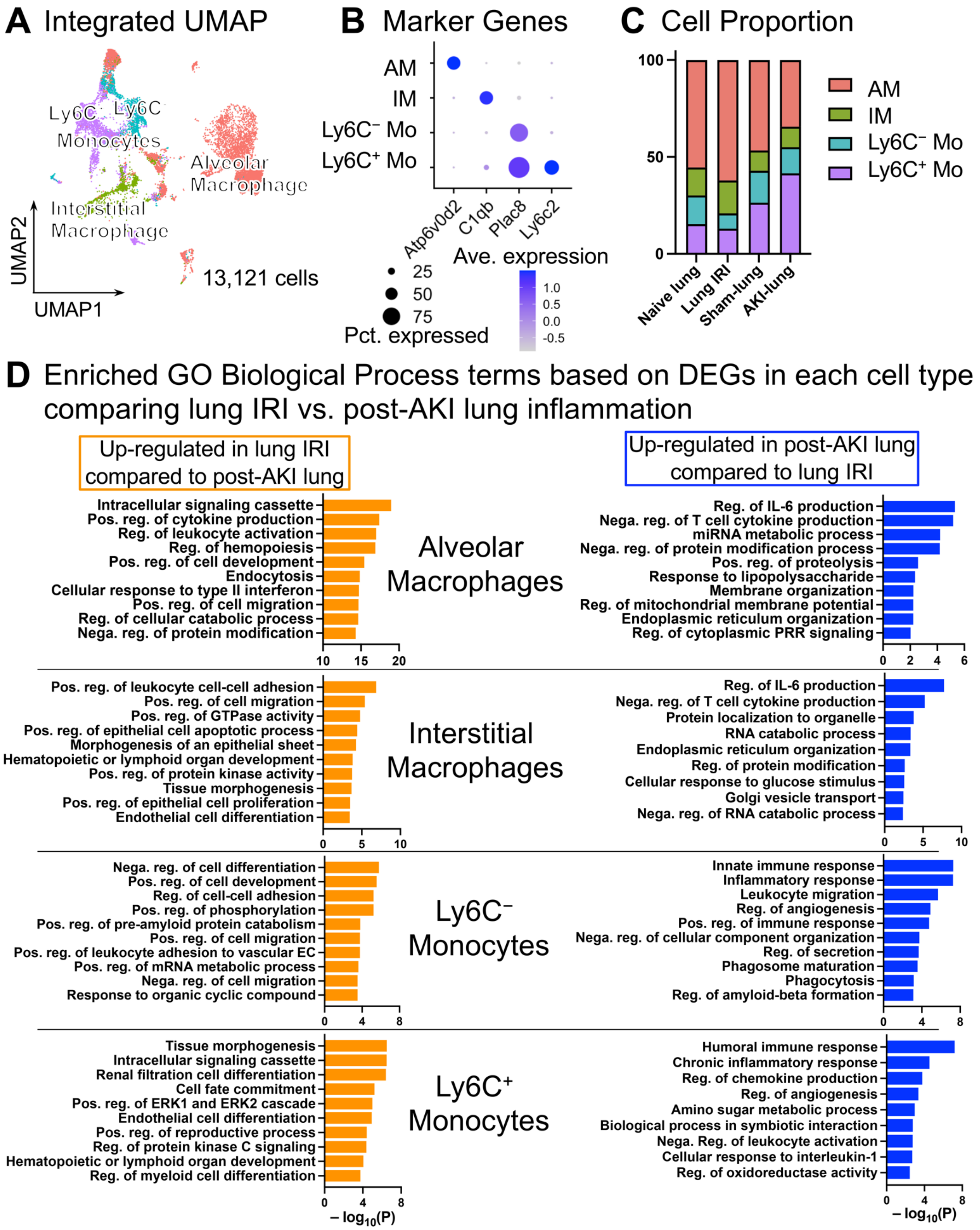


**Supplemental Figure 7. Comprehensive transcriptomic comparison of monocyte/macrophage populations in direct and indirect lung inflammation.**

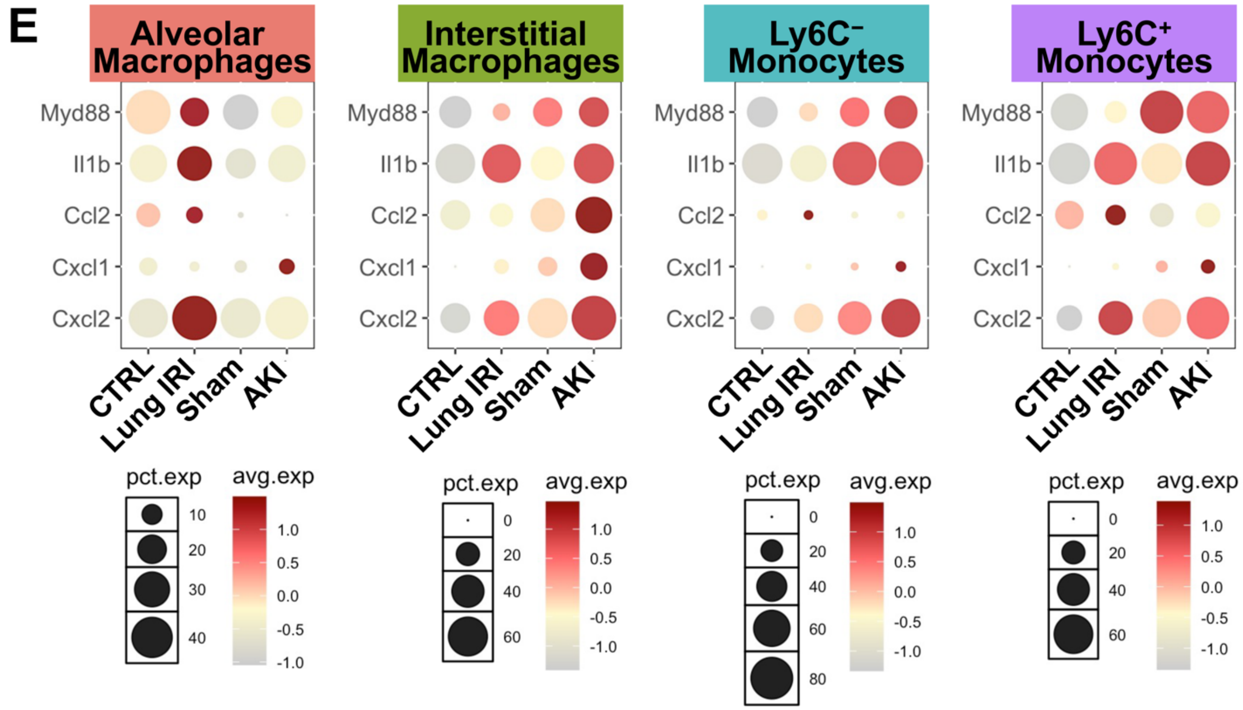


**Supplemental Figure 7. Comprehensive transcriptomic comparison of monocyte/macrophage populations in direct and indirect lung inflammation** *(continued)***.** (**A**) Syngeneic lung transplantation was used as a model of direct lung injury, as it recapitulates lung warm ischemia-reperfusion injury (IRI). After CCA integration, clustering, and annotation of post-transplant and post-AKI scRNA-seq datasets, 13,121 cells from monocyte/macrophage populations were analyzed. (**B**) Expression of marker genes for each cell type. (**C**) Proportion of each cell type across all conditions. Alveolar macrophages expanded after lung IRI, while an increase in Ly6C$^{+}$ monocytes was prominent in post-AKI lung. (**D**) Enriched Gene Ontology (GO) Biological Process terms based on differentially expressed genes (DEGs) in each cell type comparing lung IRI (direct lung injury) vs. post-AKI lung inflammation (remote lung inflammation). Several terms associated with acute inflammation such as "cytokine production", "leukocyte activation", and "cell migration" were overrepresented in alveolar macrophages after direct lung IRI. Notably, these terms were also found in intravascular (Ly6C$^{-}$ and Ly6C$^{+}$) monocytes in post-AKI lung inflammation. (**E**) Expression of inflammatory molecules in monocyte/macrophage populations. The dot size represents the percentage of cells expressing each gene, and the color scale reflects average gene expression levels.

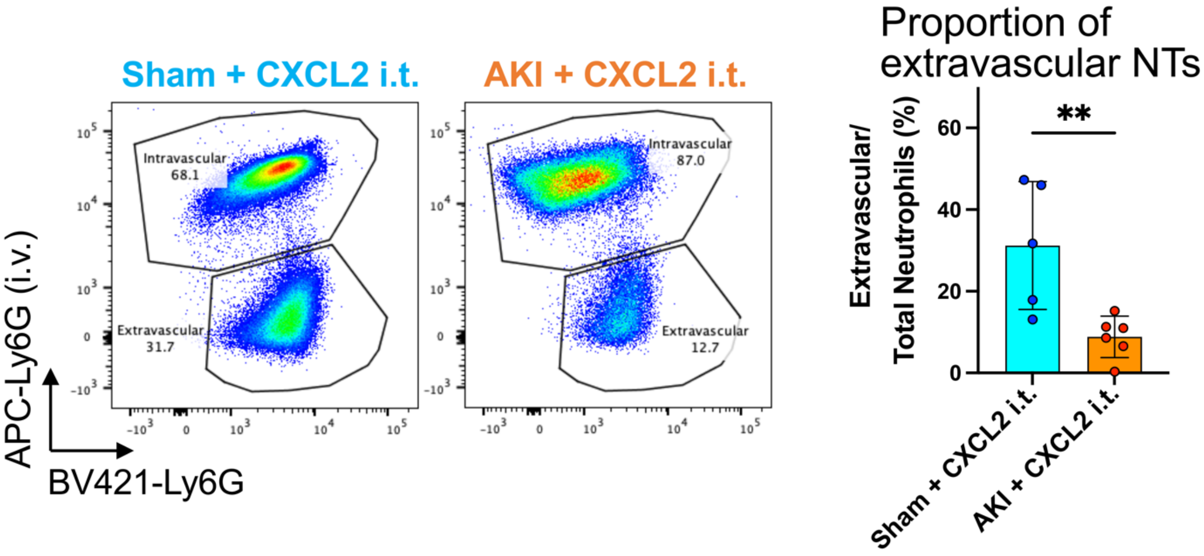


**Supplemental Figure 8. Lung neutrophil (NT) extravasation induced by intratracheal instillation of CXCL2 after sham or AKI surgery.** FACS quantification of extravasated neutrophils 24 hours after sham or AKI surgery. Mouse CXCL2 protein (0.01 µg/g) in 50 µL of sterile saline was administered intratracheally immediately after surgery. The proportion of extravasated neutrophils relative to the total neutrophil population was quantified. Data represent mean ± SD and n = 5–6 mice per group. **P < 0.01
